# Transcriptional regulator PAX4 links Receptor Tyrosine Kinases (RTKs) and cytoskeleton stability in Alzheimer’s disease and type 2 diabetes

**DOI:** 10.1101/2020.01.26.920512

**Authors:** Piyali Majumder, Kaushik Chanda, Debajyoti Das, Brijesh Kumar Singh, Partha Chakrabarti, Nihar Ranjan Jana, Debashis Mukhopadhyay

**Author notes:** Corresponding Author: Debashis Mukhopadhyay, Ph.D., Professor, Biophysics and Structural Genomics Division, Saha Institute of Nuclear Physics, HBNI, Block-AF, Sector 1, Bidhannagar, Kolkata, WB, 700064, India.

## Abstract

Alzheimer’s Disease (AD) and Type 2 Diabetes (T2D) share a common hallmark of insulin resistance. Besides Insulin Receptor (IR), two non-canonical RTKs, ALK and RYK, exhibit significant and consistent functional downregulation in *post-mortem* AD and T2D tissues. Incidentally, both have Grb2 as a common downstream adapter and NOX4 as a common ROS producing factor. Here we show that Grb2 and NOX4 play critical roles in reducing the severity of both the diseases. The study demonstrates that the abundance of Grb2 in degenerative conditions, in conjunction with NOX4, reverse cytoskeletal degradation by counterbalancing the network of small GTPases. PAX4, a transcription factor for both Grb2 and NOX4, emerges as the key link between the common pathways of AD and T2D. Both ALK and RYK downregulation elevate the PAX4 level by reducing its suppressor ARX via Wnt/β-Catenin signaling pathway. For the first time, this study brings together RTKs other than Insulin Receptor (IR), their common transcription factor PAX4 and both AD and T2D pathologies on a common regulatory platform.

## Introduction

Epidemiological studies show that type 2 diabetes (T2D) increases the risk of Alzheimer’s disease (AD) by at least 2-fold, although there are only a few mechanistic studies that provide a clear pathophysiological link (1). PET and MRI studies show marked impairment of glucose and energy metabolism in both T2D and AD (2) with amyloidogenesis being a salient feature in both. Reportedly, diabetic mice overexpressing islet amyloid polypeptide (IAPP) develop oligomers and fibrils with more severe diabetic trait akin to AD mice models overexpressing amyloid precursor protein (APP) (3). Additionally, traits like insulin resistance, altered amyloid metabolism, synaptic dysfunction, activation of the inflammatory response pathways and impairment of autophagy have been shown as common pathological features in both the diseases (4). With the understanding that several pathological signals are being shared by AD and T2D, AD has been suggested to be a neuroendocrine disorder resembling T2D (5). Insulin receptor (IR), a receptor tyrosine kinase (RTK), further links both of them exhibiting resistance to insulin signaling and other metabolic disbalances. Our recent study lists the differential activities of several human RTKs in post-mortem brain tissues of AD patients and liver tissues of T2D patients and categorizes them into functional and regulational clusters (6). Two RTKs, Anaplastic lymphoma kinase (ALK) and receptor-like tyrosine kinase (RYK), functionally behave in a similar fashion in both the disease situations (7). Both ALK and RYK are involved in the regulation of Wnt/β-Catenin signaling (8–10), which behaves aberrantly in both AD and T2D (11). In a recent study both ALK and RYK were shown to have significantly decreased activities in AD and T2D conditions (7).

Besides these pathways and signaling modalities, ROS mediated oxidative stress has been shown to be common in both the diseases (12). A ROS producing NADPH oxidase, NOX4, is constitutively active with its regulatory protein tyrosine kinase substrate (Tsk4/5) with multiple SH3 domains (13) and interacts with Grb2 naturally (14, 15). Reportedly, in AD *post-mortem* brain, in AD cell models as well as AD mouse model (APP/PS1 mouse) Grb2 transcript expression shows significant upregulation (16). In AD brain, the NOX4 expression is significantly increased with substantial correlation between its activity and age-dependent increases of Aβ and cognitive dysfunction (17, 18). In T2D, pancreatic β-cell dysfunction is promulgated by NOX4 overactivity and is sufficient to induce insulin resistance (19). Growing number of evidence suggest that Nox4-derived ROS contributes to oxidative stress during the initial and chronic stages of T2D (20).

In recent times the therapeutic potential of a cell-lineage specific transcription factor PAX4 is under scanner for T2D treatment (21). In T2D, adult β-cell mass is prevented to re-enter the cell cycle through cyclin D1 and D2 or both. Forkhead transcription factor, FoxO1, interacts with PDX-1 to modulate β-cell proliferation (22). Insulin inhibits FoxO1 through Akt-mediated phosphorylation and nuclear exclusion, which in turn increases the PDX1 expression and β-cell proliferation (23). Here, PAX4 regulates the development of islets of Langerhans by increasing the β and δ-cells population (24). Concomitantly, another transcription factor, Aristaless Related Homeobox (ARX), upregulates the α-cell population while reducing the β and δ-cells proliferation (25). Furthermore, PAX4 and ARX are not only antagonistic functionally, they negatively regulate the transcriptions of their own genes (25, 26). The transcription factor PAX4 has been shown to be involved in inducing regenerative capacity in insulin-positive islet cells in mice (21, 27). *PAX4* mutation have also been implicated in T2D (28) and its overexpression leads to β-cell proliferation and reduces apoptosis (29). Only very recently *PAX4* has been implicated in denervation in mice (30) and neurodegeneration, especially Parkinson’s disease (31). Reportedly, ARX mutation is associated with neurodegenerative and neurodevelopmental disorders through the impairment of the Wnt/β-Catenin signaling pathway (32–34).

Hippocampal and select cortical neurons in AD manifest phenotypic changes indicative of neurons re-entering cell division cycle (35, 36). Pathological signals trigger the Grb2/SOS/Ras cascade (37) that initiates cell cycle re-entry and proliferation but remains incomplete due to the lack of cell division machinery. Grb2 happens to be an important adapter of the insulin receptor and interestingly, insulin resistance is a noteworthy phenotype common to AD and T2D.

With this backdrop, the present study focuses on the regulation of both Grb2 and NOX4 by transcription factors like PAX4 and non-canonical RTKs in the context of AD and T2D. Consequences of pathological perturbations in both the diseases, starting from RTK signaling through their downstream effectors, have been tracked to link them with the cytoskeleton stability and the underpinning transcriptional regulation. Specifically, the study attempt to interpret the effects of contrasting and extraneous signals, akin to AD and T2D, that utilize similar signaling gateways to achieve common outcome in two apparently diverse diseases.

## Materials and Methods

### AD and T2D human tissues

AD (NB820-59363) and Non-AD (NB820-59177) post-mortem whole brain lysates and Type 2 Diabetic (NB820-59232) and Non-Diabetic (NB820-59291) post-mortem whole liver lysates were purchased from Novus Biologicals. For statistical reasons we procured products from different patients with different lot numbers (for patients details see Table S1a and Table S1b).

### Ethics Statement

All animal experimentations using AD and T2D mouse models were conducted following the institutional guidelines for the use and care of animals and approved by the Institutional Animal and Ethics Committee of NBRC, Gurgaon (NBRC/IAEC/2012/71) and IICB, Kolkata (IICB/AEC/Meeting/2016/AUG), respectively.

### AD and T2D mouse model

APP/PS1 or B6C3-Tg(/APPswe,PSEN1dE9/)85Dbo/J mice were obtained from the Jackson Laboratory (USA) and maintained. These transgenic mouse line for AD expresses human APPswe mutations (K670N and M671L) and exon 9-deleted human presenilin 1(PSEN1dE9) under the control of the mouse prion gene promoter. Animals were provided water and food /ad libitum/. The genotyping was carried out using PCR as described previously (38). The T2D model C57BL/6 mice 8–10 weeks of age were separated in two groups for normal chow and high-fat diet (HFD) containing 45% fat and 5.81 kcal/g diet energy content (# 960192; MP Biomedicals).

### AD and T2D cell models, Cell culture, transfection, Plasmids, siRNAs and Antibodies

Clones of both AICD-GFP construct or ‘AICD” and Grb2-DsRed construct or ‘Grb2’ were available in the lab (39–43). ALK (siRNA ID: s1269; no. 4392420), RYK (siRNA ID: s12390; no. 4390824) and PAX4 (Assay ID s10061, 4392420) siRNAs were purchased from ambion, life technologies™. Aβ-peptide (A980), Sodium Palmitate (P9767) and Insulin (I9278-5ML) were purchased from Sigma-Aldrich. Amylin (ab142398) was purchased from Abcam. Antibodies were purchased from abcam and CST (see Table S2 for details).

Human neuroblastoma (SHSY-5Y) and liver carcinoma (HepG2) cell-lines were obtained from National Cell Science Centre, Pune, India and were cultured in respective media of DMEM-F12 (Gibco) and DMEM (Gibco) supplemented with 10% fetal bovine serum (Gibco) at 37°C in 5% CO2 atmosphere under humidified condition. Transfection of cells were done using Lipofectamine 2000 (Invitrogen) and as described [23]. For co-transfection, constructs were taken in equal proportions. Human neuroblastoma (SHSY-5Y) were transiently transfected with AICD and were externally treated with 0.5μM Aβ 1-42 (Sigma A980) for 3 hours after transfection and samples were collected after 48 hours of addition (44). 0.75 mM aqueous Sodium Palmitate (Sigma P9767) along with 1% fatty acid free BSA (Sigma A8806-5G) was added to 24 hours serum starved media of HepG2 cells. Aqueous Amylin (Abcam ab142398) was added to the media at a final concentration of 0.5μM after 3 hours treatment of Palmitate. Samples were then collected after 16 hours with or without insulin (Sigma I9278-5ML) (100 nM) shock of 10 minutes.

### Protein from mammalian cell

Phosphate buffer saline (PBS) washed pellet from cell lines were lysed on ice in lysis buffer (1M Tris-HCl, pH 7.5, 1N NaCl, 0.5 M EDTA, 1M NaF, 1M Na_3_VO_4_, 10% SDS, 20mM PMSF, 10% Triton X-100, 50% glycerol) for 30 min in presence of complete protease inhibitor (Roche Diagnostics) and centrifuged at 13,000 g for 15 min. Protein concentration was determined by Bradford protein estimation assay. Western blots and quantification were done as per described protocol (16).

### Fluorescence-activated cell sorting (FACS) and estimation of ROS activity

Palmitate/ Amylin treated HepG2 cells were harvested and stained with CM-H_2_DCFDA (5-(and-6)-chloromethyl-2′,7′ dichlorodihydrofluoresceindiacetate, acetyl ester) according to manufacture’s protocol. The cells were then analysed for ROS activity by FACS ( BD FACS Calibur platform, USA).

### RNA isolation, c-DNA preparations and real time PCR

RNA was isolated from cells by TRIzol Reagent (Invitrogen,USA) extraction method following manufacturer’s protocol which discussed in Majumder et al, 2017 (16). Real time RT-PCR reaction was carried out using Syber green 2X Universal PCR Master Mix (Applied Biosystems, USA) in ABI Prism 7500 sequence detection system. The absolute quantification given by the software was in terms of CT values. The relative quantification of target genes was obtained by normalizing with internal control gene (GAPDH gene). Primer sequences and PCR conditions are mentioned in Table S3.

### Chromatin Immuno Precipitation (ChIP)

We used High Sensitivity ChIP Kit (ab185913) to perform the ChIP assay and followed manufacturer’s protocol. qRT-PCR analysis was done with the purified DNA using primers for *GRB2* and *NOX4* gene, more specifically around the *PAX4* binding sequence. Primer details are given in Table S4.

### Statistical Analysis

Unpaired ‘t’ test was carried out to compare the means of two experimental groups. The error bar represents standard error [(standard deviation/ √n); n= sample size]. To arrive at the statistically significant sample size for each experiment we did power analysis using the *a priori* model (45) as incorporated in the G*power 3.1 (46) software (16).

## Results

### Expression of Grb2 and NOX4 increases in disease models and ALK/RYK doubleknock-down condition

Having known that Grb2 and NOX4 were common at the downstream of both AD and T2D, their transcriptional and translational levels were measured. The expression level of NOX4 was elevated significantly by 1.86 fold [Fig. 1A a and b] in AD *post-mortem* brain as opposed to non-AD control. Grb2 and NOX4’s expression levels were higher by 5.27 and 2.4 folds in human *post-mortem* liver tissue of T2D patients with respect to control, respectively [Fig. 1C]. Similarly, the transcript levels of Grb2 in T2D mice were raised by 3.01 fold [Fig. 1D] and that of NOX4 in both AD and T2D mice showed significant upregulation of 5.06 and 1.45 folds, respectively [Fig. 1B and E]. NOX4 showed significant upregulation in T2D [Fig. S1] cell models as well.

**Figure 1:**
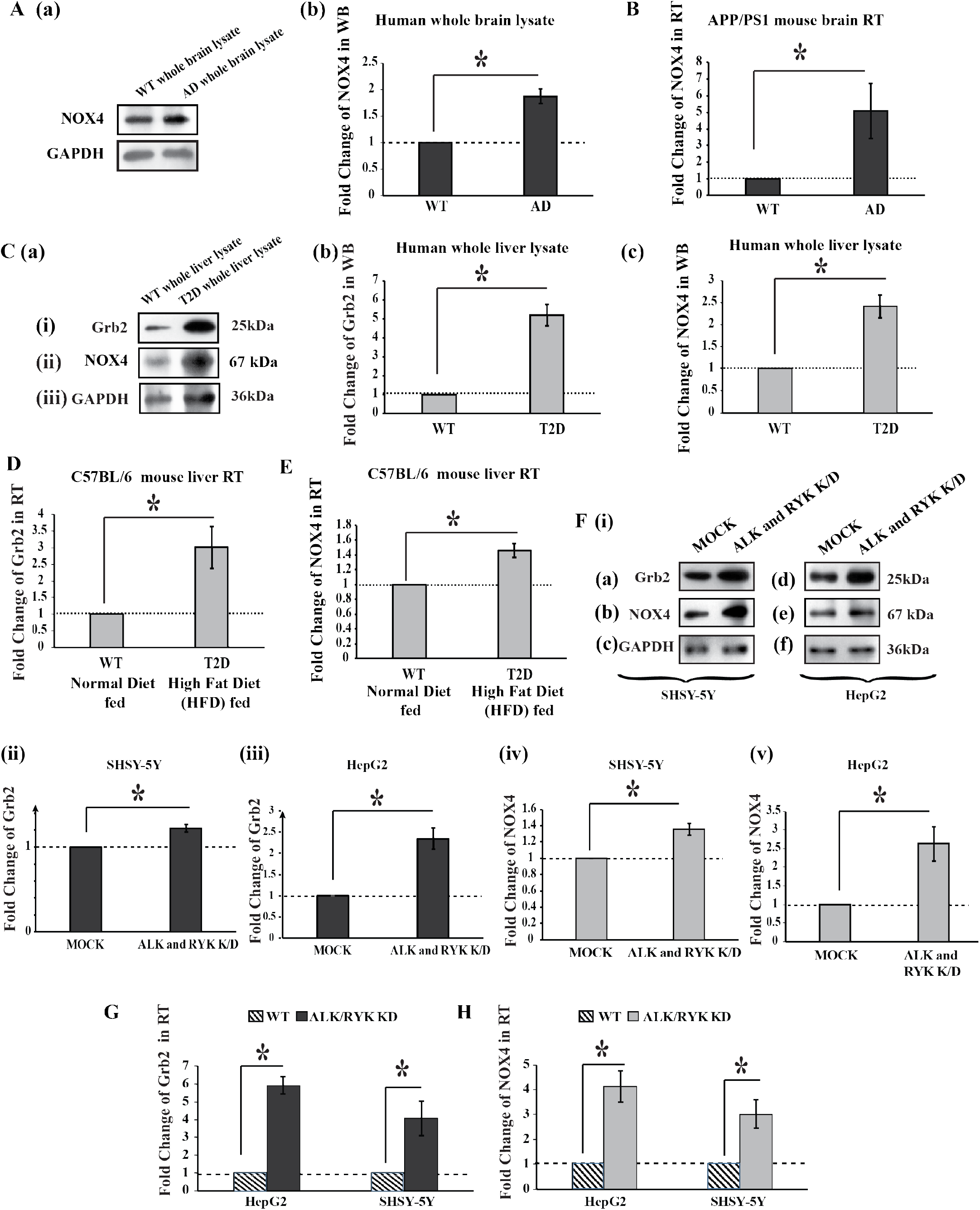
Alteration of expression of Grb2 and NOX4 in clinical, in AD or T2D mimicking mouse and in ALK and RYK double knockdown models. **A (a)** Western blot showing the NOX4 and GAPDH levels in AD whole brain lysate**. (b)** Histogram representing the mean value of optical density of the bands, normalized against GAPDH. NOX4 was overexpressed by 1.86 fold in AD Brains. **B** Shows NOX4 transcript level being upregulated by 5.06 fold by qRT-PCR of APP/PS1 mouse brain tissue. **C (a)** Western blot showing the Grb2, NOX4 and GAPDH levels in T2D whole liver lysate**. (b) and (c)** Histograms representing the mean values of Grb2 and NOX4 normalized against GAPDH. Grb2 and NOX4 were overexpressed in T2D liver by 5.27 fold and 2.4 fold, respectively. **D and E** show qRT-PCR results in T2D mouse model where C57BL/6 mice are maintained in HFD, Grb2 and NOX4 transcript levels are overexpressed by 3.01 fold and 1.45 fold, respectively. **F (i)** Western blot showing levels of Grb2 **(a and d)**, NOX4 **(b and e)** and GAPDH **(c and f)** in ALK and RYK double knock-down SHSY-5Y and HepG2 cells. **F (ii), (iii), (iv) and (v)** graphically represent the normalized overexpressed levels of Grb2 and NOX4 compared to respective controls in SHSY-5Y and HepG2 cell lines. Grb2 significantly upregulated by 1.22 fold in SHSY-5Y and 2.34 fold for HepG2 and NOX4 is increased by 1.35 fold for SHSY-5Y and by 2.62 fold for HepG2. **G and H** show transcript level changes in ALK and RYK double knockdown (K/D) conditions. Transcript levels of Grb2 significantly upregulated by 4.08 fold for SHSY-5Y and 5.9 fold in HepG2, NOX4 transcript levels increased by 3.02 fold for SHSY-5Y and by 4.13 fold for HepG2. All the statistical information is available on supplementary Table S7.

Double knockdown models for both ALK and RYK genes in SHSY-5Y and HepG2 cells [Fig. S3 and Supplementary Materials S1] were designed. Grb2 protein and transcript levels showed significant upregulation for double knockdown condition of SHSY-5Y (1.22 fold in protein level 4.08 fold transcript level) [Fig. 1 F (i)(a), (ii) and G] and HepG2 cell lines (2.34 fold in protein level 5.9 fold transcript level), respectively [Fig. 1 F (i)(d), (iii) and G]. Similarly, protein and mRNA expression levels for NOX4 was also elevated significantly for both SHSY-5Y (1.35 fold in protein level 3.02 fold transcript level) [Fig. 1 F (i)(b), (iv) and H] and HepG2 cell lines (2.62 fold in protein level 4.13 fold transcript level) [Fig. 1 F (i)(e), (v) and H]. The results resembled the conditions in diseased tissues.

### PAX4 regulates the transcription of *GRB2* and *NOX4*

While investigating for the molecular players behind *GRB2* and *NOX4* transcriptional upregulation, we used Transfac^®^ MATCH1.0 http://www.gene-regulation.com/pub/programs.html#match) online search tool to identify transcription factor binding sites upto 10Kb upstream of both *GRB2* and *NOX4* genes. In case of *GRB2*, 36 probable binding sites for 18 different transcription factors [Table S5] were found, whereas, for *NOX4*, 32 probable binding sites for 16 different transcription factors [Table S6] were found. Comparing the results for both *GRB2* and *NOX4*, three (Nkx2-5, Foxd3 and PAX4) top hits were selected among the six common (Nkx2-5, Foxd3, PAX4, CHOP C/EBP, Oct1 and Evi-1) transcription factors. The transcript levels of these three transcription factors were measured for both AD and T2D cell models [Fig. 2 A (i) and (ii)] and it was seen that whereas the mRNA levels of *Nkx2-5* were downregulated those of *PAX4* and *FOXD3* were significantly upregulated in both the models. One could summarize from quantitative RT-PCR data that *PAX4* showed maximum upregulation in both AD and T2D models where both *GRB2* and *NOX4* showed elevated expressions. PAX4 was thus selected as the major player behind altered expression of *GRB2* and *NOX4*. PAX4 expression was measured by western blot and was found to be overexpressed in both AD and T2D cell models compared to controls [Fig. S2]. This was validated with clinical samples of AD and T2D [Fig. 2 B], where PAX4 expression was upregulated by 1.8 and 2.00 folds respectively. Consequences of PAX4 knockdown, using Silencer^®^ select *PAX4* siRNA construct, and significantly reducing the endogenous *PAX4* level in SHSY-5Y and HepG2 cell lines [Fig. 2 C (iii)], have been measured. Although non-significant in SHSY-5Y, the transcript level of *GRB2* decreased significantly in HepG2 [Fig. 2 C (i)]. PAX4 knockdown on the other hand significantly decreased the level of *NOX4* in SHSY-5Y but in case of HepG2, the increase was insignificant [Fig. 2 C (ii)].

**Figure 2:**
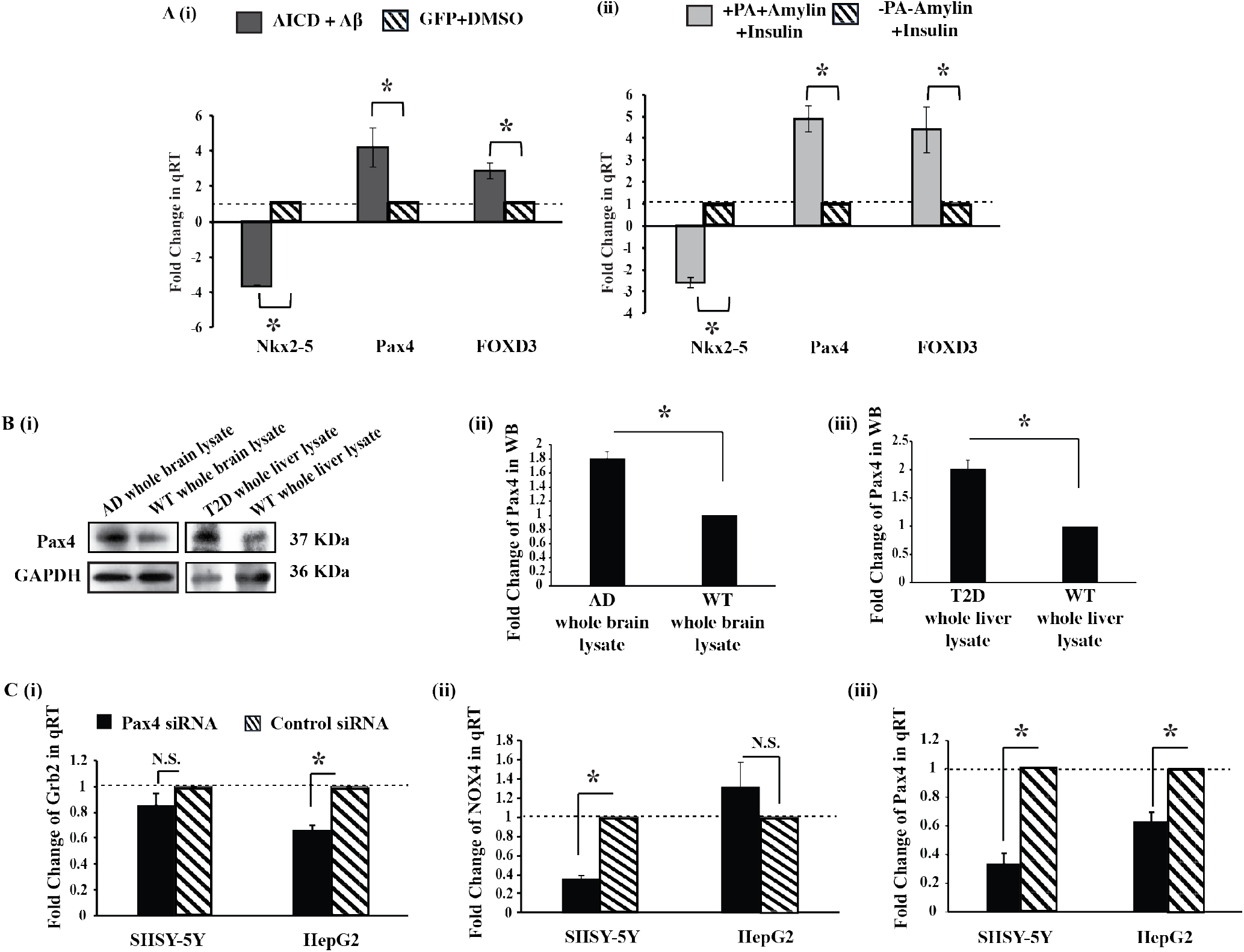
Role of PAX4 in transcription of Grb2 and NOX4. **A (i)** shows the transcript level alterations of −3.6 fold for *Nkx2-5*, +4.19 fold for *PAX4* and +2.89 fold for *FOXD3* in AD (AICD+Aβ) cell models compared to control. Similarly, **(ii)** shows the transcript level alterations of −2.59 fold for *Nkx2-5*, +4.88 fold for *PAX4* and +4.39 fold for *FOXD3* in T2D (+PA+Amylin+Insulin) cell models compared to controls. **B** shows PAX4 protein overexpression by 1.8 and 2.00 folds by Western Blot in **(ii)** AD patients’ whole brain samples and **(iii)** T2D patients’ whole liver samples, respectively. **C** displays the alterations in transcript levels of **(i)** *GRB2* [in SHSY-5Y 1.16 fold and in HepG2 1.52 fold], *NOX4* [in SHSY5Y 2.83 fold and in HepG2 2.3 fold] and **(iii)** endogenous *PAX4* [in SHSY5Y 2.99 fold and in HepG2 1.58 fold] in PAX4 knockdown situation in SHSY5Y and HepG2 cell lines. All the statistical information is available on supplementary Table S7.

Considering a possible correlation between downregulation of the expressions and activities of ALK and RYK and upregulation of the transcription factors, the mRNA levels of *Nkx2-5, PAX4* and *FOXD3* have been measured in *ALK*/*RYK* double knockdown cells and qRT-PCR results showed +5.75, +5.84 and +3.13 folds’ alterations, respectively, in SHSY-5Y [Fig. 3 A (i)] and +2.91, +4.8 and +2.46 fold changes, respectively, in HepG2 [Fig. 3 A (ii)]. Resembling AD and T2D cell models, in *ALK*/*RYK* double knockdown cells *PAX4* showed maximum upregulation among the transcription factors. Interestingly, both the knockdown (ALK/RYK) conditions showed significant upregulation of PAX4 protein levels (1.65 and 1.98 folds, respectively, for SHSY-5Y and HepG2 cell lines) [Fig. 3 B].

**Figure 3:**
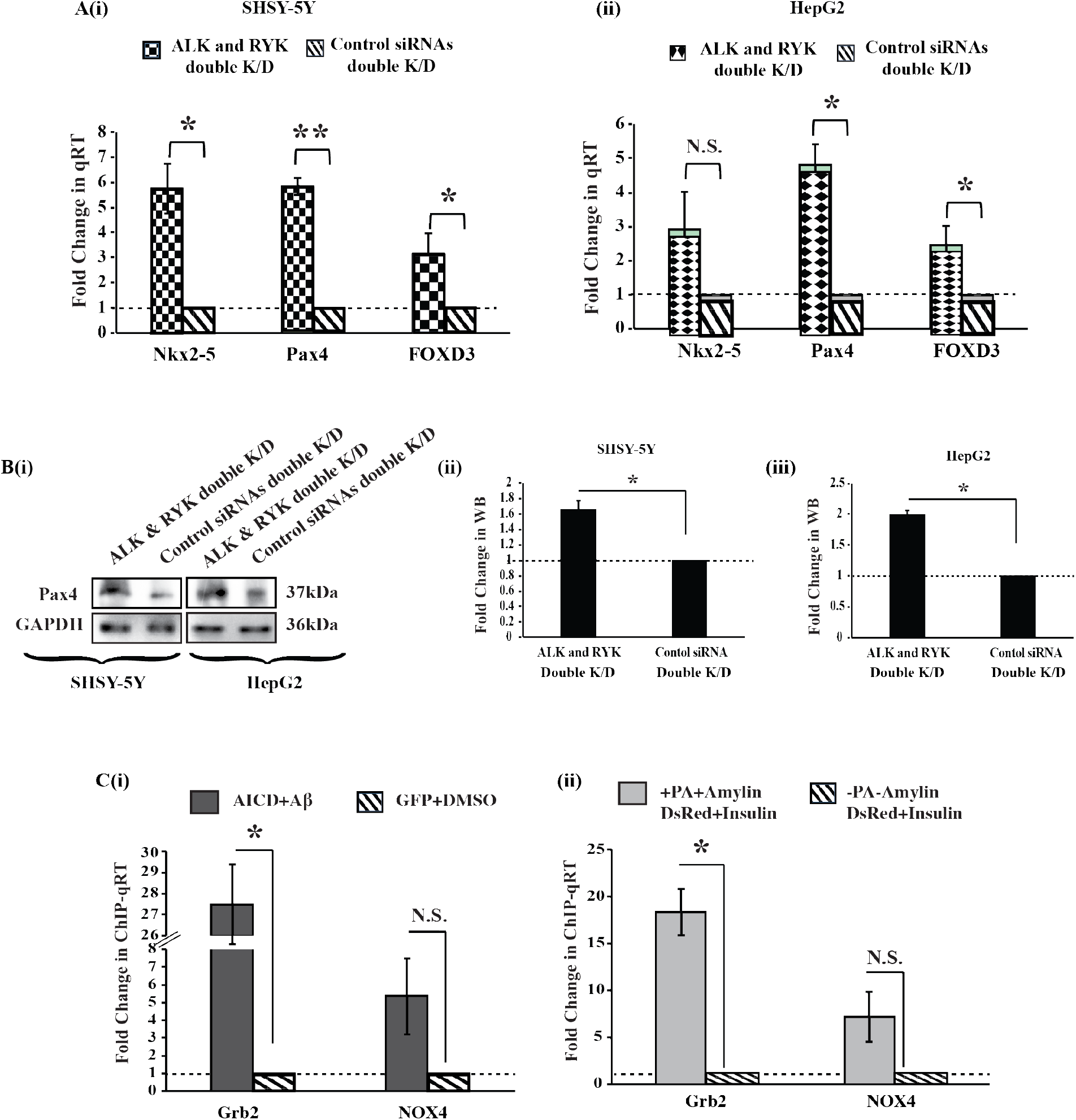
ALK and RYK double knockdown effect on PAX4 levels. **A** shows the transcript level alterations of *Nkx2-5, PAX4* and *FOXD3* in ALK and RYK double knockdown (K/D) situation in **(i)** SHSY-5Y [+ 5.75 fold for *Nkx2-5*, +5.84 fold for *PAX4* and +3.13 fold for *FOXD3*] and **(ii)** HepG2 cell line [+2.91 fold for *Nkx2-5*, +4.81 fold for *PAX4* and +2.46 fold for *FOXD3*]. **B (i)** shows the PAX4 protein level alterations by Western Blot in ALK and RYK double knockdown (K/D) model in both SHSY-5Y and HepG2 cell lines. **(ii) and (iii)** histograms that graphically denotes the upregulation of endogenous PAX4 levels in double knockdown model in both SHSY-5Y [+1.65 fold] and HepG2 cells [+1.98 folds]. **PAX4 binds to the upstream region of Grb2 and NOX4 gene**. **C** shows ChIP data where it proves that PAX4 significantly upregulates *GRB2* expression in both **(i)** AD [+27.49 fold] and **(ii)** T2D [+18.33 fold] cell models by binding at its upstream region and acting as an enhancer like transcription factor. In case of *NOX4*, PAX4 fails to significantly upregulate its expression in both (i) AD [5.35 fold] and (ii) T2D [7.22 fold] cell models. All the statistical information is available on supplementary Table S7.

Chromatin Immuno-precipitation (ChIP) assays confirmed elevated levels of direct recruitment of PAX4 protein in the *GRB2* and *NOX4* upstream regions. For the AD cell model PAX4 recruitment increased by 27.49 and 5.35 folds [Fig. 3 C (i)] and in T2D cell model they were elevated by 18.33 and 7.22 folds [Fig. 3 C (ii)], for *GRB2* and *NOX4* genes, respectively.

### ARX correlates the ALK/RYK downregulation with PAX4 via β-Catenin signaling

In the course of exploring the mechanism behind PAX4 upregulation in the degenerative diseases, the role of ARX has emerged out in a significant way. The significant upregulation of ARX transcript level by 3.3 and 3.1 folds in PAX4 knockdown conditions in both SHSY-5Y and HepG2 cells established the mutually repressive natures of ARX and PAX4 [Fig. 4 A]. Moreover, the ARX transcript levels were significantly downregulated by 2.6, 3.7, 3 and 3.7 folds in the AD and T2D cell models and *ALK/RYK* double knockdown situation of both SHSY-5Y and HepG2 cells, respectively [Fig. 4 B]. Additionally, the β-Catenin’s role as ARX-regulator was also established by measuring its expression levels. The Western blots showed significant downregulation of the β-Catenin by 3.5 and 1.5 folds in AD and T2D cell models, respectively [Fig. 4 C]. The *ALK/RYK* double knockdown also showed a significant β-Catenin downregulation of 1.15 and 6.5 folds in both SHSY-5Y and HepG2 cells, respectively [Fig. 4 D].

**Figure 4:**
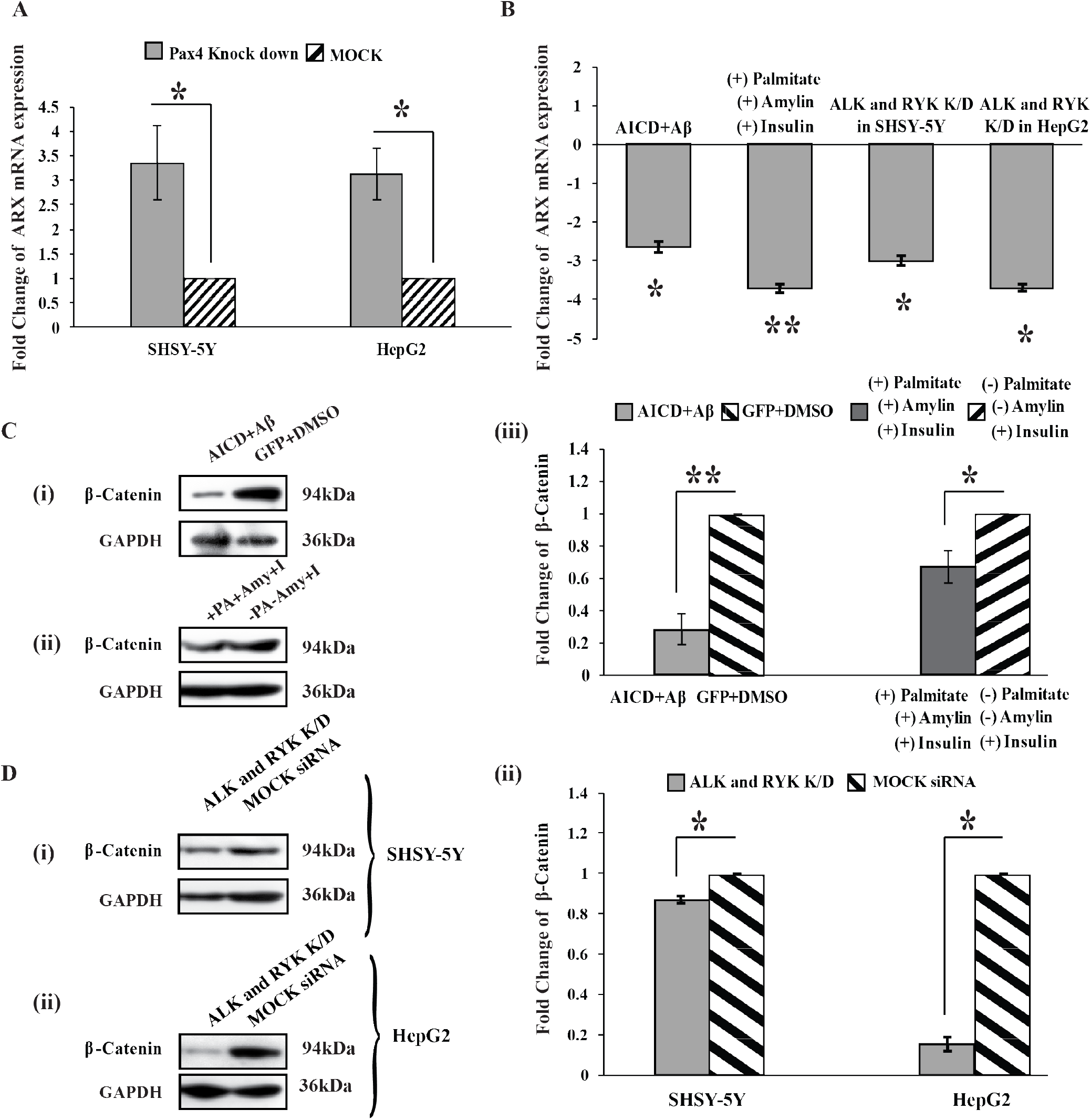
Role of ARX and β-Catenin in AD and T2D cell model and in ALK/RYK double knockdown model. **(A)** shows that ARX transcript level is being upregulated in PAX4 knockdown condition by 3.35 and 3.13 fold by qRT-PCR in both SHSY-5Y and HepG2 cells, respectively. **(B)** shows the downregulation of ARX transcript levels by 2.6, 3.7, 3 and 3.7 folds in AD (AICD+Aβ) and T2D (+PA+Amylin+Insulin) cell models and also in ALK/RYK double knockdown condition in SHSY-5Y and HepG2 cells, respectively. **(C)** Western blot showing the reduction of β-Catenin expression levels in **(i)** AD and **(ii)** T2D cell model. **(iii)** Histogram representing β-Catenin being downregulated by 3.5 and 1.5 fold in AD and T2D cell model, respectively. **(D)** shows the downregulation of β-Catenin expression levels in ALK and RYK double knockdown situation in both (i) SHSY-5Y and (ii) HepG2 cells by 1.15 and 6.5 folds, respectively. All the statistical information is available on supplementary Table S7.

### Expressions of cytoskeletal proteins change in conjunction with ALK/RYK deactivation

Considering significant cytoskeletal degradation in AD like scenario (16), the protein and transcript levels of four cytoskeleton proteins (viz., α-Tubulin, Vimentin, α-Smooth muscle actin (α-SMA) and Stathmin1) were compared between whole liver tissue of T2D patients and non-Diabetic whole liver samples. The relative protein levels were downregulated by 1.16 fold for α-Tubulin, 1.83 fold for Vimentin, 1.68 fold for α-SMA and 1.95 fold for Stathmin1, respectively [Fig. 5 a]. The transcript levels also showed significant (−5.3 fold α-tubulin; −6.49 fold Vimentin; −2 fold α-SMA and −2.9 fold Stathmin1) downregulation in T2D mice [Fig. 5 B]. Similarly, in the T2D cell model the relative protein levels were downregulated by 2.7 fold for α-Tubulin, 1.98 fold for Vimentin, 1.27 fold for α-SMA and 1.97 fold for Stathmin1, respectively [Fig. 5 C]. Continuing with the attempt to link degeneration with ALK/RYK deactivation, we compared the protein and the transcript levels of α-Tubulin in *ALK*/*RYK* double knockdown conditions. A significant reduction in protein levels (2.64 and 1.42 fold, respectively) was observed in both SHSY-5Y and HepG2 cell lines [Fig. 5 D]. Similarly, transcript levels of α-Tubulin were also downregulated by 2.3 and 1.5 folds for both SHSY-5Y and HepG2 cell lines, respectively [Fig. 5 E]. In both Palmitate and Amylin treated situations, the percentage recovery of cytoskeletal proteins from disassembly upon Grb2 overexpression upregulated the expressions of α-Tubulin, Vimentin, α-SMA and Sthathmin1 by 98.2%, 84.39%, 508.8% and 62.2% respectively [Fig. 5 F].

**Figure 5:**
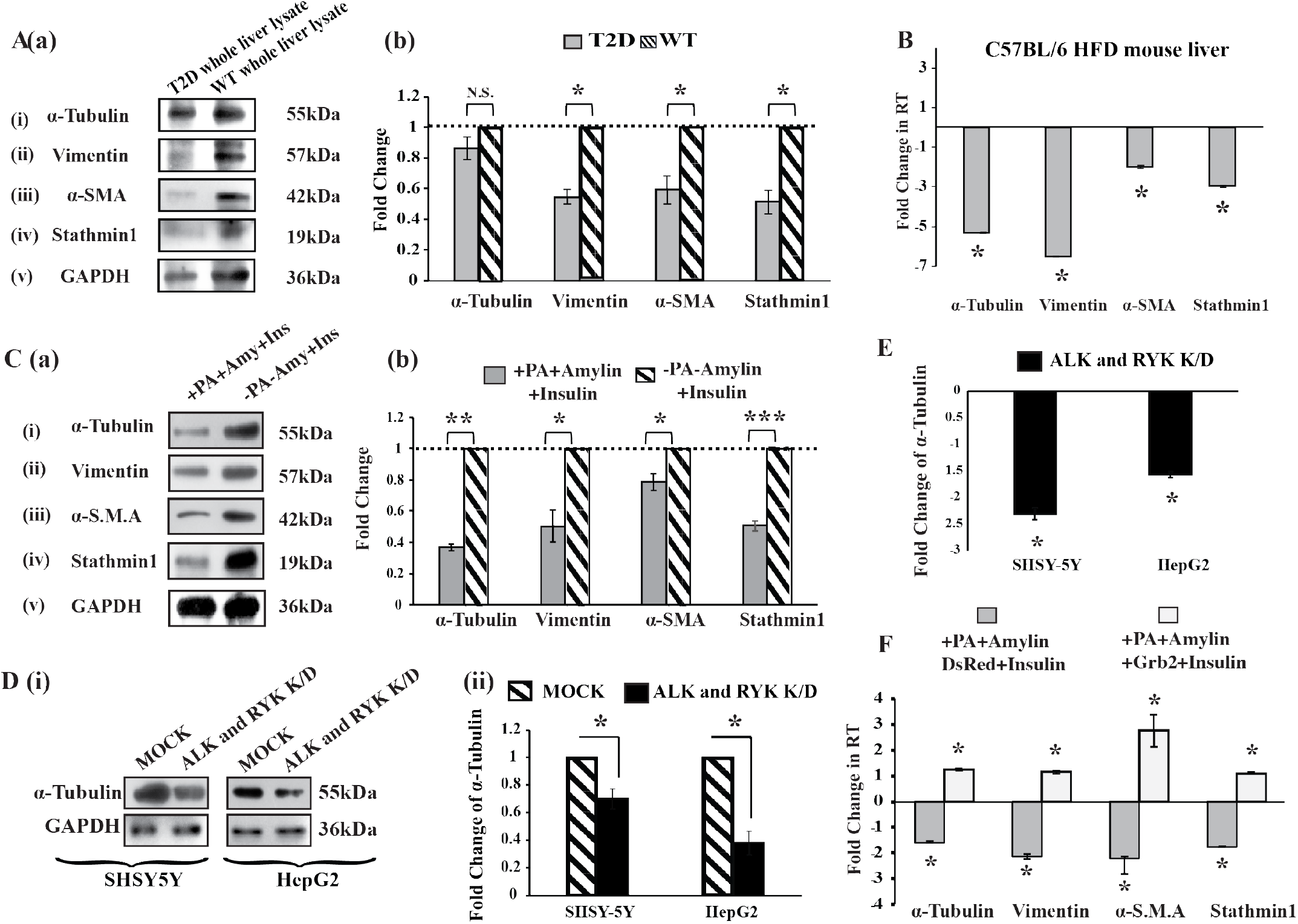
Participation of ALK and RYK in signaling events leading to cytoskeleton degradation and Grb2 mediated reversal in T2D: A. cytoskeleton degradation. **a** Representative western blots (n=2) of four cytoskeletal proteins (i) α-Tubulin, (ii) Vimentin, (iii) α-SMA and (iv) Stathmin1 with (v) GAPDH used as internal control in human diabetic whole liver lysates, compared to wild type whole liver lysate. **b** Histogram representing the mean value of optical density of the protein bands, normalized against GAPDH, with a decrease of 1.16 fold for α-Tubulin, 1.83 fold for Vimentin, 1.68 fold for α-SMA and 1.95 fold for Stathmin1. Samples are derived from the same experiments and the blots are processed in parallel. **B** Shows transcript level changes in in α-Tubulin (−5.3 fold), Vimentin (−6.49 fold), α-SMA (−2 fold), and Stathmin1 (−2.9 fold) by qRT-PCR in HFD fed C57BL/6 mice compared to normal diet fed controls. **C a** Western blots depict alteration in the expression of **(i)** α-Tubulin, **(ii)** Vimentin, **(iii)** α-Smooth muscle actin (α-SMA) and **(iv)** Stathmin1with GAPDH in T2D cell model (PA+Amylin+Insulin induced HepG2 cells). The samples are derived from the same experiments and the blots are processed in parallel. **b** Histograms showing changes in the four cytoskeletal proteins, normalized by GAPDH, with decrease 2.7 fold for α-Tubulin, 1.98 for Vimentin, 1.27 fold for α-SMA and 1.97 fold for Stathmin1**. D ALK and RYK double knockdown condition controls α-Tubulin degradation. D(i)** shows by western blot, the decrease of α-Tubulin in both SHSY-5Y [−2.64 fold] and HepG2 cell line [−1.42 fold] in ALK and RYK double knockdown (K/D) situation. **E** shows the α-Tubulin transcript level downregulation in ALK and RYK double knockdown (K/D) model in both SHSY5Y [−2.3 fold] and HepG2 cell lines [−1.5 fold] by qRT-PCR. **F** Shows transcript level changes for the four cytoskeletal proteins in T2D (PA+Amylin+Insulin induced condition) with or without Grb2. All the statistical information is available on supplementary Table S7.

### The signaling pathways also show a reversal of outcome through Grb2-NOX4 interaction

While investigating the pathways that could be responsible for degradation of cytoskeletal proteins, involvement of three small GTPases viz., RhoA, Rac1 and Cdc42 [Fig. 6 A (a) (i), (ii), (ii) and (b)] was observed. Similar to AD situations, under Palmitate and Amylin treated disease inducing conditions, the expression levels of RhoA and Rac1 decreased significantly by 2.13 (n=3) and 1.63 folds (n=3) respectively and the activity of Cdc42 increased by 64.2% (n=3) folds. Down the line, the significantly reduced Cofilin activity bounced back by 43.2 % [Fig. 6 B (a) (i) and (b) (i)] upon Grb2 overexpression and Palmitate/Amylin treatment. Besides, SSH-1’s (which is also an activator of Cofilin through dephosphorylation) distal upstream effector NOX4 (47–49) was overexpressed significantly by 1.31 fold post Palmitate/Amylin treatment that did reduce significantly by 1.26 fold [Fig. 6 B(a) (ii) and (b) (ii)] upon Grb2 overexpression. Considering that α-Tubulin was downregulated in T2D conditions, the activity of Gsk3β was also analyzed [Fig. 6 C]. The activity of Gsk3β, a kinase that would hyperphosphorylate Tau leading to destabilization of the microtubule network, was found to increase significantly by 1.7 fold [Fig. 6 C (a) (i) and (b)] under disease inducing Palmitate/Amylin treated condition. The activity of its upstream effector AKT1 decreased significantly by 1.54 fold [Fig. 6 C (a) (ii) and (b)] under T2D inducing condition. Interestingly, Nox4, an interactor of Grb2, was found to be 1.34 fold overexpressed endogenously in T2D cell model but reduced in the presence of Grb2 [Fig. 6 B(a) (ii) and (b) (ii)]. The interaction of Grb2 with NOX4 enhanced 1.35 fold in T2D scenario [Fig. 7 A (a)]. Further, the intensity of interaction of NOX4 and Grb2 was checked in T2D mouse liver lysate that increased 2.87 fold in T2D situation compared to the wild type [Fig. 7 A (b)]. In addition to this reversal, ROS activities of Palmitate/Amylin and Palmitate/Amylin/Grb2 were measured by flow cytometry by using CMH2-DCFDA ROS indicator. Although overexpressed Grb2 could significantly reverse the effects of many T2D related metabolic changes, to our surprise in the working T2D like model, over-expression of Grb2 not only failed to reduce the activity level of Reactive Oxygen Species (ROS) but rather elevated it by 3.57 fold [Fig. 7 B]. Nevertheless, extent of this limited fate reversal upon Grb2 overexpression could be estimated and in both Palmitate and Amylin treated situations, Grb2 overexpression upregulated the expressions of cytoskeletal proteins significantly.

**Figure 6:**
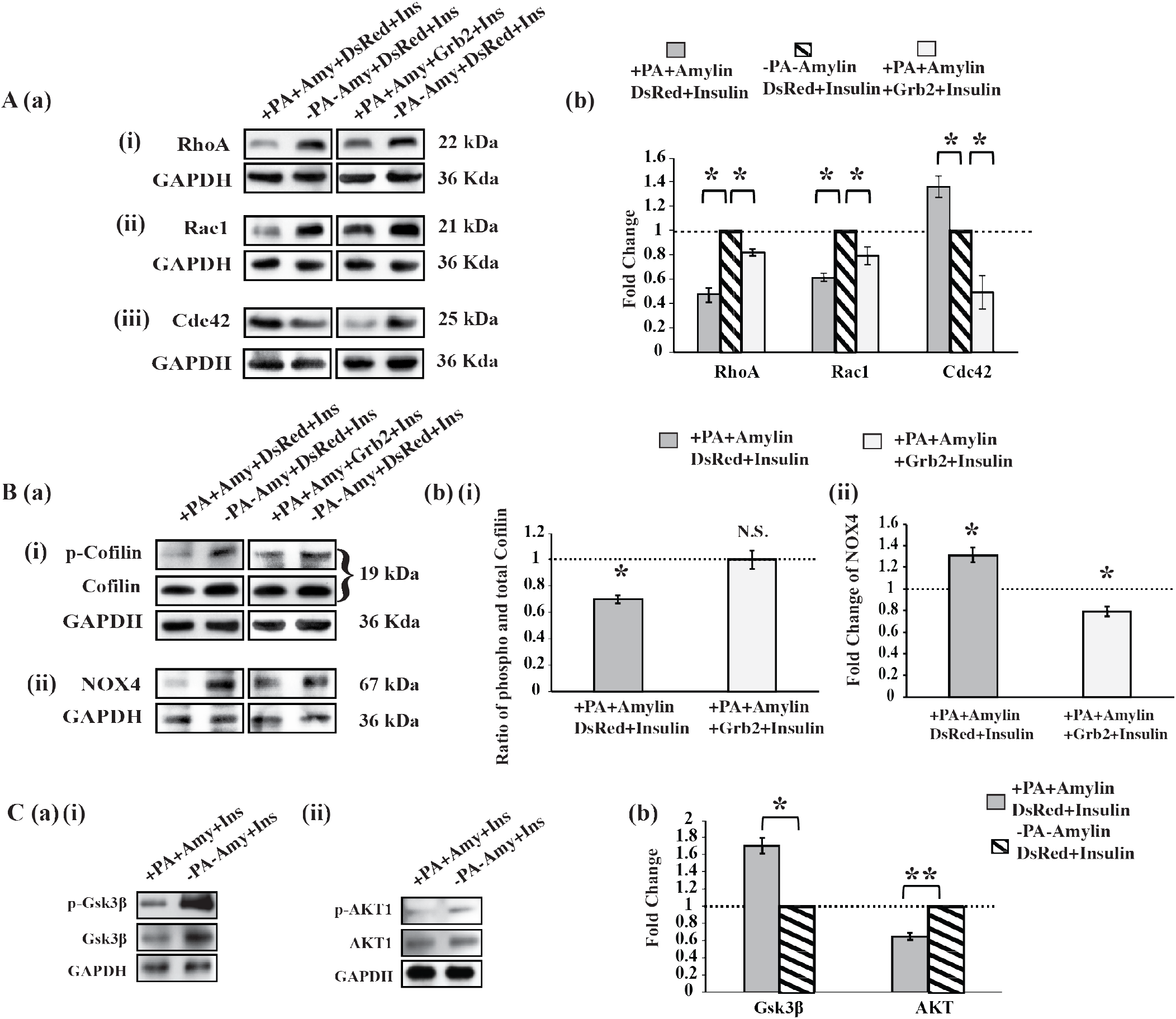
Signaling molecules participate in T2D. **A(a)** shows western blot for activity changes of small GTPases i.e., **(i)** RhoA [−2.13 fold and in presence of Grb2 −1.22 fold], **(ii)** Rac1 [−1.63 fold and in presence of Grb2 −1.26 fold] and **(iii)** CDC42 [+1.36 fold and in presence of Grb2 −2.05 fold] in T2D inducing (+PA+Amylin+Insulin) and reversing conditions (+PA+Amylin+Insulin +Grb2). In **A(b)** Bar diagram represents the activity alterations for the small GTPases. **B(a),** Depicts the protein level or activation of signaling molecules like **(i)** Cofilin [−1.43 fold and in presence of Grb2 1.000 fold] and **(ii)** NOX4 [+1.31 fold and in presence of Grb2 −1.26 fold] respectively, by western blot. **B(b) (i) and (ii)** show graphical representation of the alteration of Cofilin and NOX4 respectively. **C(i)** shows the activity alteration of GSK3β [+1.7 fold] and AKT1 [−1.54 fold] in T2D inducing (+PA+Amylin+Insulin) condition compared to wild type condition (-PA-Amylin+Insulin) and **(ii)** shows the histogram of activity alteration of both GSK3β and AKT1. All the statistical information is available on supplementary Table S7.

**Figure 7:**
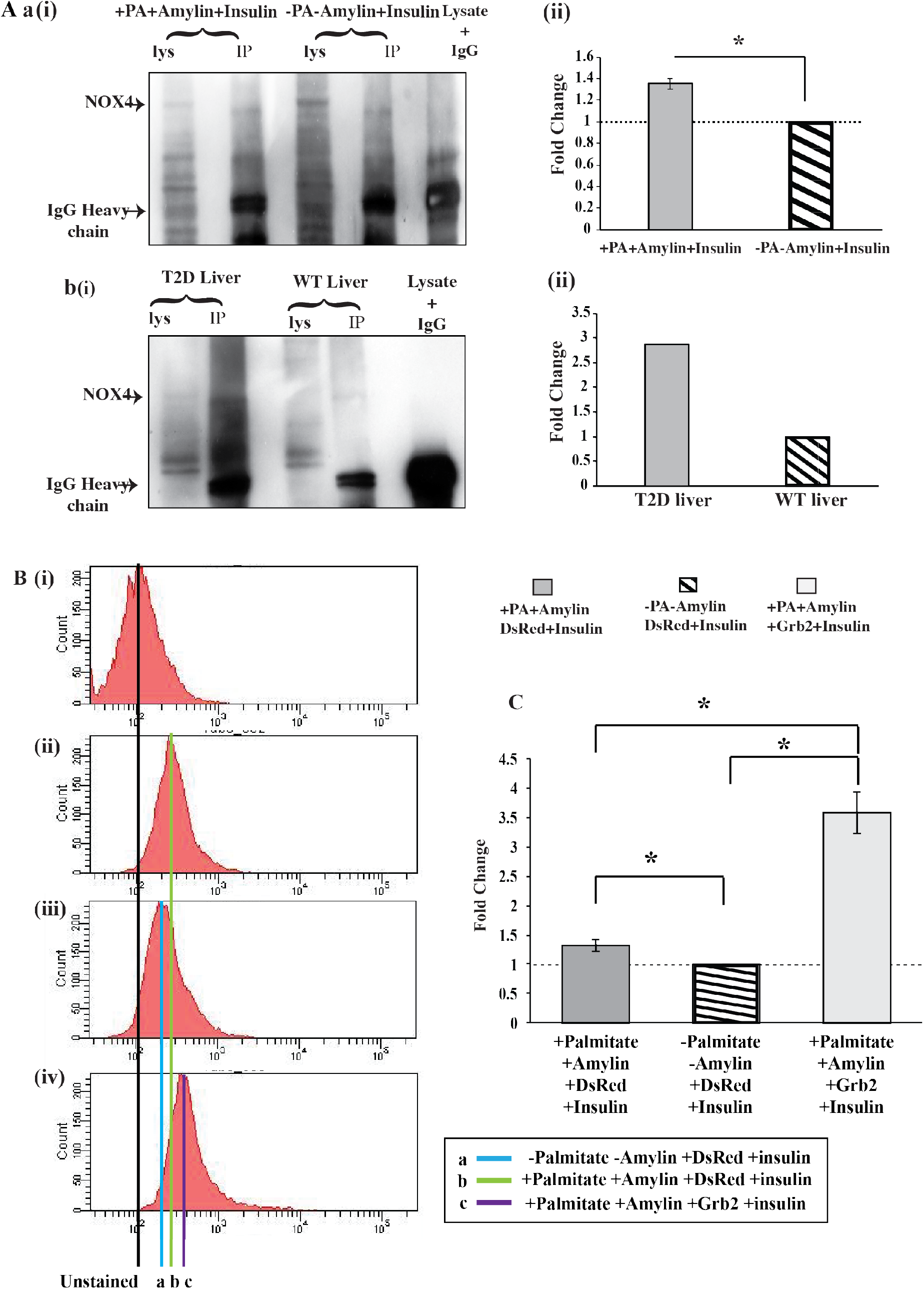
Grb2 and NOX4 interaction prevents cytoskeletal degradation in T2D like scenario and ROS activity. **A (a) (i)** shows the variation of interaction of Grb2 and NOX4 in T2D inducing (+Palmitate, +Amylin and +Insulin) conditions compared to controls (-Palmitate, -Amylin and +Insulin). **A (a) (ii)** graphical representation of the intensity of the interaction between NOX4 and Grb2. **A (b) (i)** shows the variation of interaction of Grb2 and NOX4 in type 2 diabetic mouse (HFD C57BL/ 6) whole liver lysate compared to WT (Normal fed C57BL/6) control. **A (b) (ii)** graphical representation of the intensity of the interaction between NOX4 and Grb2 in T2D mouse liver lysate. **ROS activity Assay. B (i), (ii), (iii) and (iv)** show the FACS data for ROS activity of unstained, -Palmitate -Amylin +DsRed +Insulin, +Palmitate +Amylin +DsRed +Insulin and +Palmitate +Amylin +Grb2 +Insulin, respectively. **C** Shows graphically that ROS activity increases in T2D inducing (+Palmitate +Amylin +DsRed +Insulin) condition by 1.3 fold and in presence of Grb2 ROS activity further significantly increases by 3.5 fold. All the statistical information is available on supplementary Table S7.

## Discussion

This work unveils the commonality of two diverse diseases, AD and T2D. Over a decade, continuous efforts has been taken to establish a relationship between the two, but almost none of them focused on the signaling components and their possible convergence on a single transcription factor PAX4.

The precise common signaling mechanism for AD and T2D is still unclear, albeit the fact that both the diseases confer insulin resistance. Up until our recent work, Insulin Receptor (IR) was the only RTK known to be involved in the signaling of both diseases. Profiling the activities of other RTKs in the two diseases opened up a larger picture and the toxic effects of both Aβ and Amylin oligomers were predicted to have impacts on other receptor tyrosine kinases (7). Amongst all the common RTKs, ALK and RYK were the two showing similar levels of deactivation in all working models of AD and T2D (7). ALK, a member of insulin receptor superfamily, is commonly known for its involvement in many cancer types, especially in non-small-cell lung cancer (NSCLC) (50). RYK is a co-receptor of non-canonical Wnt signaling (51). Grb2 happens to be a common downstream adapter for both (52). It was shown previously that Grb2 strengthened its interaction with NOX4 to rescue the cytoskeleton degradation in AD like situations (16). We postulated that the two RTKs, ALK and RYK, might act like non-canonical receptors with the potential to become a significant link between AD and T2D via Grb2 and NOX4.

The downstream consequences at the molecular level, for both the signals, were significant compromise of cytoskeleton integrity. Interestingly, ALK and RYK double knockdown cell lines also resembled similar phenotypes [Fig. 5 D and E]. Scanning the signaling pathways that could connect the common RTKs with downstream cytoskeleton degradation, we singled out Grb2 and NOX4, both of which had elevated levels in the disease scenarios [Fig. 1]. Several reports on Grb2 strongly suggested its role in cellular survival in AD (41, 43). Transcript levels of the cytoskeletal protein components were checked and were found to increase several folds with Grb2’s overexpression in both AD (16) and T2D models [Fig. 5 F].

It was prudent therefore to try to understand the underpinning cellular mechanisms that help in the cytoskeletal restructuring, with possibly Grb2 playing a pivotal role. Activities of three small GTPases, RhoA, Rac1 and Cdc42, known to regulate cell morphology through rearrangement of cytoskeletal proteins (53–55) acting as molecular switches, were significantly altered and again renormalized with overexpression of Grb2. These alterations in the activities of small GTPases were sufficient to perturb the downstream signaling events and, ultimately, the cytoskeletal proteins degraded primarily through Cofilin mediated pathways. Cofilin is one of the downstream effectors of LIMK1 whose de-phosphorylation enables actin severing and depolymerizing activities (56). These results convincingly implicated similar pathways for the rearrangement of the cytoskeleton network in AD and T2D. Grb2’s intervention in both disease models significantly inactivates Cofilin by phosphorylation (16) [Fig. 6 B (a) (i), (b) (i)], a crucial regulator of actin Dynamics (57). Similar to the case in AD, in reversing the signaling pathways involved in the degradation of cytoskeletal proteins, Grb2 was also involved in T2D as well. Abnormal NOX activation was reported in T2D (58). NOX4 was found to interact with Grb2, activating Src (15).

Adapter protein NOX4, from ROS production pathway, could also regulate Cofilin activity through Slingshot homolog-1 (SSH-1), the phosphatase of Cofilin. Both the disease models showed significant upregulation of NOX4, which was subsequently decreased with Grb2’s overexpression [Fig. 6 B(a)(ii) and (b) (ii)] (16). These small checks and balances, at different levels along the signaling cascades, culminated in a larger scale perturbation in the cytoskeleton network and Grb2’s role emerged out as that of reversal of these perturbations to some extent [Fig. 5 F] (16). Furthermore, while examining other effects on cytoskeleton integrity due to Grb2’s overexpression, it was noted that NOX4 interacted with Grb2 in normal conditions (14, 15) which increased several folds under both AD (16) and T2D disease conditions [Fig. 7 A]. As NOX4 was responsible for ROS generation, we also checked whether Grb2 overexpression could put a check on the ROS, which it could not. On the contrary, the ROS activity had increased in the presence of too much of Grb2 [Fig. 7 B]. This might be a hint towards the fact that even though the cells geared up the protective mechanisms, the defense was lost with the progression of the disease and beyond a threshold.

Eventually, the study unraveled the relation of ALK and RYK in controlling the cytoskeleton integrity through the overexpression of Grb2 and NOX4. Employing bioinformatics tools and quantitative real-time PCR, transcription factor, PAX4 was identified as a key regulator [Fig. 2 A]. PAX4 is thought to be one of the crucial transcription factors that regulates the gene network governing β-cell mass expansion and survival under the pathophysiological conditions of T2D (59). Recent studies have shown association of *PAX4* mutations with T2D in Japanese and Afro-Americans population (28, 60). However, *PAX4* gene dysfunction would increase the susceptibility of apoptosis along with reducing cell proliferation leading to a gradual loss of β-cell and ultimately to diabetes (59). Interestingly, the PI3K inhibitor, Wortmannin, showed potential to induce both insulin resistance and PAX4 expression (59, 61). Naturally, *PAX4* emerged as a survival gene because of its role in regulating both β-cell mass expansion and Grb2 expression. On the other hand, the mutual antagonism between PAX4 and ARX [Fig. 4 A] connects the aberrant Wnt/β-Catening signaling with the upregulation of PAX4 by the reduction of ARX in both the diseases [Fig. 4 B]. Nevertheless, this revelation assists to connect the dots between the deactivation of ALK and RYK with the downregulation of β-Catenin expression [Fig. 4 C,D] level, followed by a decrease of ARX. Thus suggesting that the downregulation of ALK and RYK elevates the PAX4 level [Fig. 2 B and 3 B], which in turn upregulates Grb2 and NOX4 in both AD and T2D.

Additionally, NOX4 could induce mitochondrial dysfunction by inhibiting the mitochondrial chain complex 1 (62). Alterations in the expression of genes coding for mitochondrial and cytoskeletal proteins contributed to the mitochondrial dysfunction observed in insulin-resistant conditions of both AD and T2D (63, 64). Interestingly, the majority of differentially expressed genes (DEGs) targeted by *PAX4* were commonly enriched in both oxidative phosphorylation pathway (OXPHOS) and neurodegenerative diseases (31). Besides in T2D, this work established PAX4 as a promising candidate for a significant upregulation of *GRB2* and *NOX4*, acting as the coveted ‘missing link’ between AD and T2D’s common signaling pathways [Fig. 2 B,C; Fig. 3 B,C and Fig. S2], involving ALK and RYK as upstream receptors and Grb2 and NOX4 as downstream adapters, eventually affecting the cytoskeleton.

For the first time two non-canonical receptors (ALK or RYK) have been shown to link disparate signals (AICD/Aβ and Palmitate/Amylin) leading to similar pathways (Grb2/PI3K/AKT/RhoA/ Rac1/ Cdc42/ Cofilin or NOX4/ SSH-1/Cofilin) that ultimately culminate in similar mechanistic consequences (degradation of cytoskeleton network) [Fig. 8]. This would help understand both the diseases from each others perspectives. A plethora of evidences indicate a strong link between T2D and AD. In the context of Insulin Receptor (IR) insensitivity being the pathological hallmark of both diseases, previous findings called for understanding the roles of other members of the RTK family. The present study showed that two RTKs’, ALK and RYK, downregulated in both AD and T2D, could induce common pathways involving Grb2 and NOX4 as pivotal molecular players with PAX4 as a common transcription factor regulating both. Alterations of Grb2 and NOX4 perturbed cytoskeleton integrity in both the cases. The work unravelled (a) a novel avenue to study the implications of other RTKs, besides IR, for both the diseases, (b) the unique role of Grb2 in the management of cytoskeleton degradation under AD/T2D conditions and (c) most importantly, emergence of PAX4 emeregence of as the first transcription factor that actively regulates the pathways of both AD and T2D.

**Figure 8:**
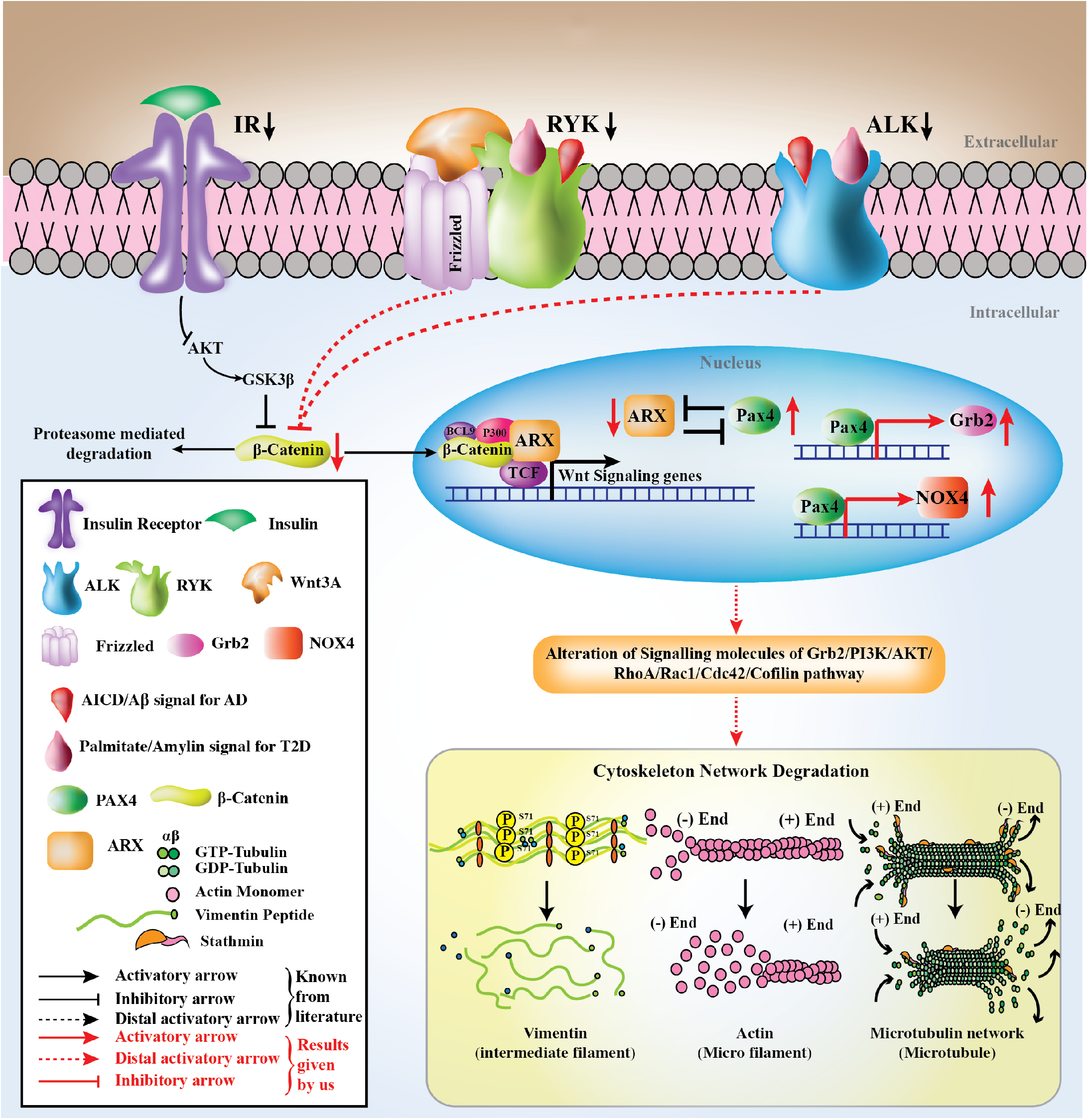
Summary Diagram. A cartoon representation Summarizing the overlapping of signaling pathways for both AD and T2D, mediated via the downregulation of membrane bound ALK/RYK, upregulation of the transcription factor PAX4 which in turn upregulates Grb2/NOX4. These effectors, through the signaling molecules, affect the fate of the cytoskeletal proteins.

## Abbreviation List

RTK: Receptor Tyrosine Kinase
ALK: Anaplastic lymphoma Kinase
RYK: Receptor related to Tyrosine Kinase
Grb2: Growth factor Receptor Bound protein 2
NOX4: NADPH Oxidase 4
PAX4: Paired Box Protein 4
ARX: Aristaless Related Homeobox
AICD: Amyloid-β Protein Precursor Intracellular Domain
Aβ: Amyloid β peptide
IR: Insulin Receptor

## Acknowledgements

We acknowledge Prof. S. Banerjee for the FACS facility and Dr. O. Chakrabarti for discussions.

## Author Contributions

- **Designed research** DM,PM
- **Performed research** PM, KC, DD, BKS
- **Contributed new reagents or analytic tools** PM, DM, PC, NRJ
- **Analyzed data** PM, DM
- **Wrote the paper** PM, DM

