## Supplementary Figures for "Transcriptional regulator PAX4 links Receptor Tyrosine Kinases (RTKs) and cytoskeleton stability in Alzheimer’s disease and type 2 diabetes"

### Figure S1

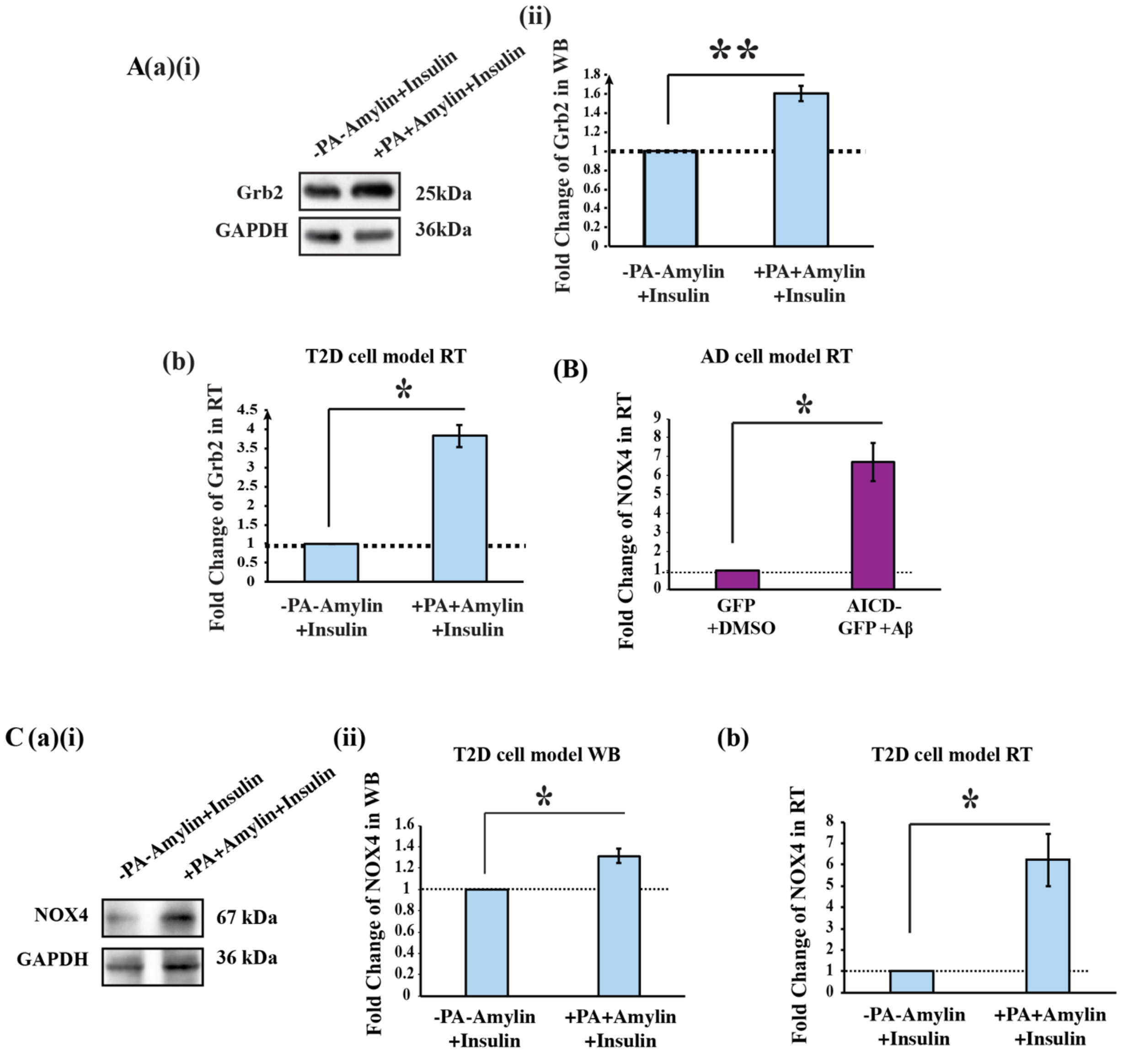


**Figure S1: Grb2 and NOX4 are upregulated in AD and T2D cell models. A (a) (i)** Western blot showing alterations of Grb2 and GAPDH levels in Palmitate and Amylin treated conditions in HepG2. **(ii)** Graphical representation of the normalized variations of Grb2 expressions compared to control cells. **(b)** Normalized fold changes of mRNA levels of Grb2 for qRT-PCR experiments with GAPDH taken as internal control. **B** shows the normalized fold changes of mRNA levels of NOX4 in AICD transfected/Aβ treated conditions in SHSY-5Ywith GAPDH taken as internal control. **C (a) (i)** Western blot showing alterations of NOX4 and GAPDH levels in Palmitate and Amylin treated conditions (T2D cell model) in HepG2. **(ii)** Graphical representation of the normalized variations of NOX4 expressions compared to control cells. **(b)** Normalized fold changes of mRNA levels of NOX4 with GAPDH taken as internal control in T2D cell model.

**Figure S2**

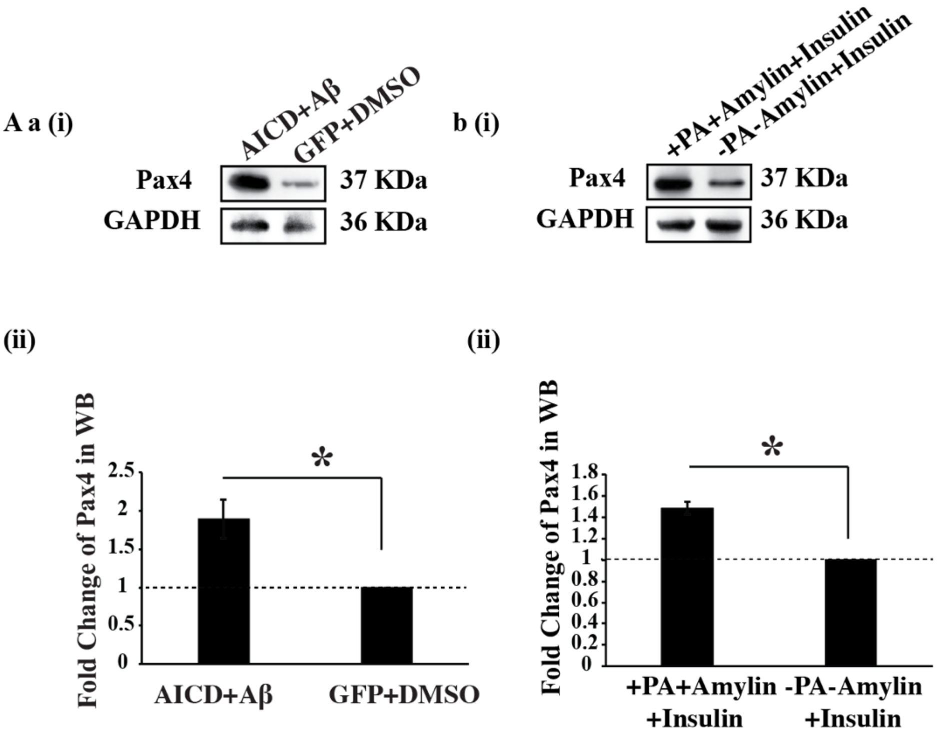


**Figure S2: Endogenous PAX4 levels are upregulated in AD and T2D cell models. A** shows PAX4 protein level alterations by Western Blot in **(a)(i)** AD cell model **(b)(i)** T2D cell model. **A (a) (ii) and (b) (ii)**graphically represent the Pax4 protein level elevation of 1.8 and 2 folds in AD and T2D models, respectively.

**Figure S3**

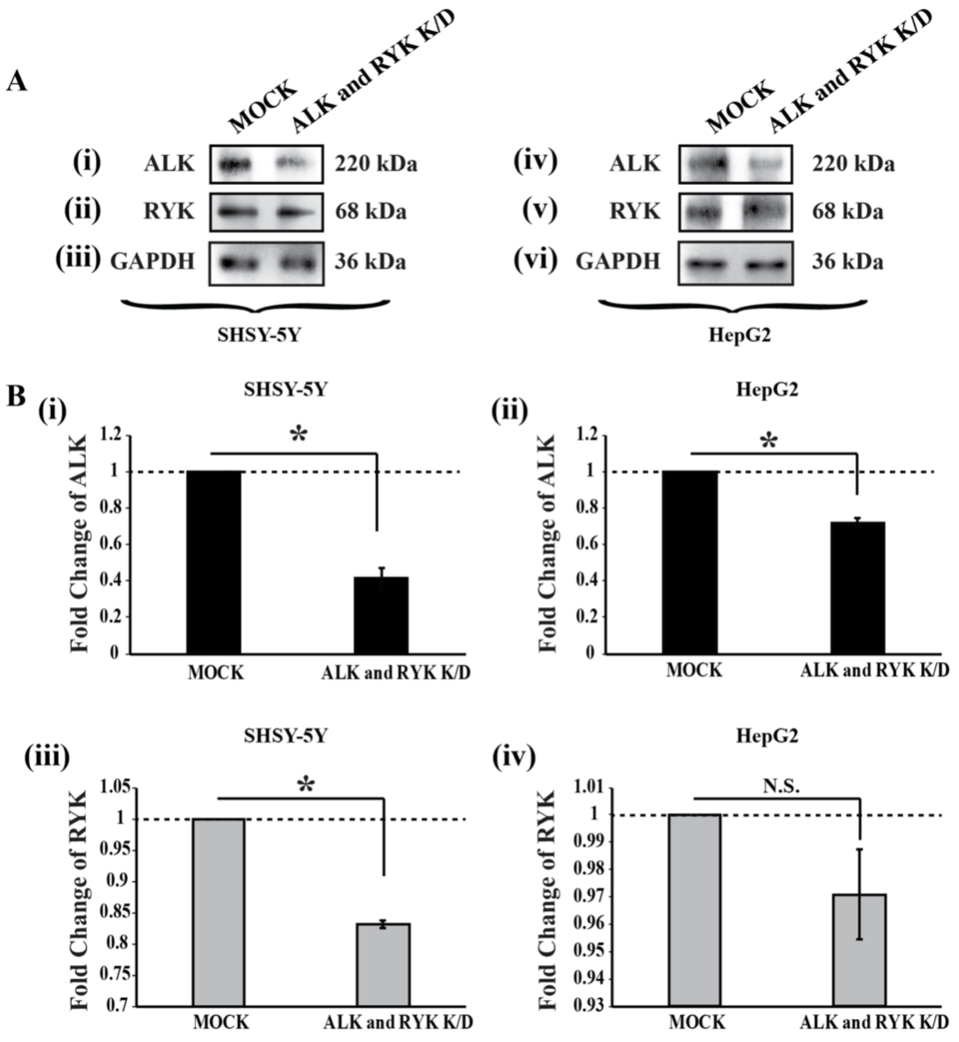


**Figure S3: Expression levels alterations of ALK and RYK in ALK/RYK double knockdown model. A** Western Blotshowing the ALK **(i and iv)**, NOX4 **(ii and v)** and GAPDH **(iii and vi)** levels in both SHSY-5Y and HepG2 cell lines. **(i) and (iii),** Graphical representations of the normalized variations of ALK and RYK in expressions compared to MOCK (two non-targeted siRNAs treated) in SHSY-5Y cell line. **(ii) and (iv)** Graphical representation of the normalized variations of ALK and RYK in expressions compared to MOCK in HepG2 cell line.
