## Supplementary Materials for "Transcriptional regulator PAX4 links Receptor Tyrosine Kinases (RTKs) and cytoskeleton stability in Alzheimer’s disease and type 2 diabetes"

**Supplementary Materials S1**

**Validation of ALK and RYK double knockdown model:** ALK and RYK double knockdown models were established in two cell lines, SHSY-5Y (human neuroblastoma cells) and HepG2 (human liver carcinoma cells). Here ALK and RYK two genes were simultaneously knocked down in cells by using siRNAs against both ALK and RYK compared to cells where two non-targeted (MOCK) siRNAs were simultaneously knocked down. The extent of knockdown was checked by measuring expression levels of both ALK and RYK by western blot in both SHSY-5Y and HepG2 cells. In SHSY-5Y cells, knockdown of ALK and RYK downregulated the expression levels of ALK and RYK by 2.42 [Fig S3 A (i) and B (i); *$p=0.0088<0.05$; $n=3$] and 1.2 folds [Fig S3 A (ii) and B (iii); *$p=0.019<0.05$; $n=2$], respectively. Similarly, in HepG2 cells, knockdown of ALK and RYK reduced the expression levels of ALK by 1.38 fold [Fig S3 A (iv) and B (ii); *$p=0.0088<0.05$; $n=2$] and RYK did not show significant decrease [Fig S3 A (v) and B (iv); N.S.; $p=0.22>0.05$; $n=2$].
