## Supplementary Tables for "Transcriptional regulator PAX4 links Receptor Tyrosine Kinases (RTKs) and cytoskeleton stability in Alzheimer’s disease and type 2 diabetes"

**Table S1: Disease Lysates' Specification list**

**Table S1a: Whole brain lysate**

| Catalog number | Lot # | Sex | Age | Ethnicity | Pathology |
|---|---|---|---|---|---|
| NB820-59177 | C101138 | M | 98 | Caucasian | Cause of death: Prostate Cancer |
| NB820-59177 | B811092 | F | 60 | Caucasian | Cause of death: unknown; had emphysema |
| NB820-59363 | B909049 | M | 72 | Caucasian | Alzheimer Disease |
| NB820-59363 | B105129 | M | 75 | Caucasian | Alzheimer Disease |

**Table S1b: Whole liver lysate**

| Catalog number | Lot # | Sex | Age | Ethnicity | Clinical diagnosis |
|---|---|---|---|---|---|
| NB820-59291 | B901104 | M | 30 | Asian | Normal |
| NB820-59291 | B901104 | M | 71 | Caucasian | Normal |
| NB820-59232 | B812314 | F | 84 | Caucasian | Type 2 Diabetes |
| NB820-59232 | C104213 | M | 78 | Caucasian | Type 2 Diabetes |

**Table S2: List of Antibodies**

| Antibody Name | Catalog Number | Dilution |
|---|---|---|
| **Grb2** | ab32037 (Abcam) | 1:5000 |
| **α-Tubulin** | ab4074 (Abcam) | 1:5000 |
| **Vimentin** | ab8069 (Abcam) | 1:5000 |
| **α-SMA** | ab32575 (Abcam) | 1:2500 |
| **Stathmin1** | ab52630 (Abcam) | 1:10,000 |
| **NOX4** | ab133303 (Abcam) | 1:1000 |
| **Slingshot (SSH-1)** | ab76943 (Abcam) | 1:1000 |
| **Phospho-Cofilin (S3)** | ab12866 (Abcam) | 1:1000 |
| **Phospho-AKT1 (S473)** | ab81283 (Abcam) | 1:1000 |
| **AKT1** | ab124341 (Abcam) | 1:1000 |
| **phospho-Gsk3β (S9)** | ab75814 (Abcam) | 1:1000 |
| **Gsk3β** | ab32391 (Abcam) | 1:1000 |
| **RYK** | ab124961 (Abcam) | 1:2500 |
| **Pax4** | ab42450 (Abcam) | 1:1000 |
| **GAPDH** | ab9484 (Abcam) | 1:5000 |
| **Cofilin** | CST-3318 | 1:1000 |
| **Phospho-ALK (Tyr1507)** | CST-14678 | 1:1000 |
| **β-Catenin** | ab224803 (Abcam) | 1:3000 |
| **ALK** | CST-3633 | 1:1000 |

**Table S3: Primer sequences and PCR conditions for qRT-PCR.**

| Name of the genes | PCR condition | PCR Cycle | Primer sequences |
|---|---|---|---|
| **α-Tubulin (mouse)** | 95°C→10min[95°C 30sce, 55°C 30 sec, 60°C 1min] 72°C→10min | 35 | Forward:5'GCAGTGTTCGTAGACCTGGAA3'<br>Reverse: 5'TTATTGGCAGCATCCTCCTT3' |
| **Vimentin (mouse)** | 95°C→10min[95°C 30sce, 55°C 30 sec, 60°C 1min] 72°C→10min | 35 | Forward:5'ATGCTTCTCTGGCACGTCTT3'<br>Reverse: 5'AGTGAGGTCAGGCTTGGAAA3' |
| **Stathmin1 (mouse)** | 95°C→10min[95°C 30sce, 55°C 30 sec, 60°C 1min] 72°C→10min | 35 | Forward:5'AGAAGGACCTTTCCCTGGAG3'<br>Reverse: 5' TTCTCATGCTCCCGCTTC3' |
| **Grb2 (mouse)** | 95°C→10min[95°C 30sce, 55°C 30 sec, 60°C 1min] 72°C→10min | 35 | Forward:5'AAATGCTCAGCAAACAGCGG3'<br>Reverse: 5'TGAAGTGCTGCACATCATTTC3' |
| **NOX4 (mouse)** | 95°C→10min[95°C 30sce, 55°C 30 sec, 60°C 1min] 72°C→10min | 35 | Forward:5'TGTTGCATGTTTCAGGTGGT 3'<br>Reverse: 5'TACTGGCCAGGTCTGCTTT 3' |
| **Pax4 (mouse)** | 95°C→10min[95°C 30sce, 55°C 30 sec, 60°C 1min] 72°C→10min | 35 | Forward:5'GGCTCCCAGTGTGTCCTCTA 3'<br>Reverse: 5'GGGACTGGGAAGAACTGGAG 3' |
| **ALK (mouse)** | 95°C→10min[95°C 30sce, 55°C 30 sec, 60°C 1min] 72°C→10min | 35 | Forward:5'CTCTGGTGCTGGTGGAGAAC 3'<br>Reverse: 5'GTCACATCGAGGAGGGACAG 3' |
| **RYK (mouse)** | 95°C→10min[95°C 30sce, 55°C 30 sec, 60°C 1min] 72°C→10min | 35 | Forward:5'GACCTGGCTGCTAGGAACTG 3'<br>Reverse: 5'CCCTAGGCAGTGGTAGTCCA 3' |
| **GAPDH (mouse)** | 95°C→10min[95°C 30sce, 55°C 30 sec, 60°C 1min] 72°C→10min | 35 | Forward:5'AGCCTCGTCCCGTAGACAAAA3'<br>Reverse: 5'TGGCAACAATCTCCACTTTGC3' |
| **α-Tubulin (human)** | 95°C→10min[95°C 30sce, 55°C 30 sec, 60°C 1min] 72°C→10min | 35 | Forward:5'CCGGGCAGTGTTTGTAGACT3'<br>Reverse: 5'GCAGCATCTTCTTTGCCTGT3' |
| **Vimentin (human)** | 95°C→10min[95°C 30sce, 55°C 30 sec, 60°C 1min] 72°C→10min | 35 | Forward:5'GGCACGTCTTGACCTTGAAC3'<br>Reverse: 5'GTGAGGTCAGGCTTGGAAAC3' |

| Name of the genes | PCR condition | PCR Cycle | Primer sequences |
|---|---|---|---|
| **α-SMA (human)** | 95°C→10min[95°C 30sce, 55°C 30 sec, 60°C 1min] 72°C→10min | 35 | Forward: 5'ACCCAGCACCATGAAGATCA3'<br>Reverse: 5'TTTGCGGTGGACAATGGAAG3' |
| **Stathmin1 (human)** | 95°C→10min[95°C 30sce, 55°C 30 sec, 60°C 1min] 72°C→10min | 35 | Forward:5' AAGGATCTTTCCCTGGAGGA 3'<br>Reverse: 5' GTTTCTCAGCCAGCTGCTTC 3' |
| **Grb2 (human)** | 95°C→10min[95°C 30sce, 55°C 30 sec, 60°C 1min] 72°C→10min | 35 | Forward:5'AGAACTGGTACAAGGCAGAGC 3'<br>Reverse: 5 'GATAAGAAAGGCCCCATCGT 3' |
| **NOX4 (human)** | 95°C→10min[95°C 30sce, 55°C 30 sec, 60°C 1min] 72°C→10min | 35 | Forward:5'CCGGCTGCATCAGTCTTAACC 3'<br>Reverse: 5 'TCGGCACAGTACAGGCACAA 3' |
| **ALK (human)** | 95°C→10min[95°C 30sce, 55°C 30 sec, 60°C 1min] 72°C→10min | 35 | Forward:5'TGGAGTTTGTCACCAGTGGA3'<br>Reverse: 5'TGTCTTCAGGCTGATGTTGC 3' |
| **RYK (human)** | 95°C→10min[95°C 30sce, 55°C 30 sec, 60°C 1min] 72°C→10min | 35 | Forward:5'GGCTGCCAGGAACTGTGT 3'<br>Reverse: 5'CCCCCAGACAGTGATAGTCC 3' |
| **Nkx2-5 (human)** | 95°C→10min[95°C 30sce, 55°C 30 sec, 60°C 1min] 72°C→10min | 35 | Forward:5'AAGAGCTGTGCGCGCTGCAGAA 3'<br>Reverse: 5'ATCTTGACCTGCGTGGACGTG 3' |
| **FOXD3 (human)** | 95°C→10min[95°C 30sce, 55°C 30 sec, 60°C 1min] 72°C→10min | 35 | Forward:5'TCTGCGAGTTCATCAGCAACG 3'<br>Reverse: 5'TTGACGAAGCAGTCGTTGAGC 3' |
| **Pax4 (human)** | 95°C→10min[95°C 30sce, 55°C 30 sec, 60°C 1min] 72°C→10min | 35 | Forward:5'GCTGAAGGGTGAGTGTCCAG 3'<br>Reverse: 5'TGGAGGAGACACTGGGAGTC 3' |
| **ARX (human)** | 95°C→10min[95°C 30sce, 55°C 30 sec, 60°C 1min] 72°C→10min | 35 | Forward:5' AGCTGTCACCCAAGGAGGAG 3'<br>Reverse: 5' GTGAAGACGTCCGGGTAGTG 3' |
| **GAPDH (human)** | 95°C→10min[95°C 30sce, 55°C 30 sec, 60°C 1min] 72°C→10min | 35 | Forward:5'TCCCTGCACCACCAACTGTTAG3'<br>Reverse: 5'GGCATGGCATGTGGTCATGAG3' |

**Table S4: Primer Sequences and PCR condition for ChIP assay**

| Name of the genes | PCR condition | PCR Cycle | Primer sequences |
|---|---|---|---|
| **Grb2 (human)** | 95$^o$C→7min[95$^o$C 10sce, 55$^o$C 10 sec, 72$^o$C 8 sec] 72$^o$C→1 min | 40 | Forward:5'TTAGGTGGTGGTGACACGCC 3'<br>Reverse: 5'ACTGTTAGTAGAGATGGGCT 3' |
| **NOX4 (human)** | 95$^o$C→7min[95$^o$C 10sce, 55$^o$C 10 sec, 72$^o$C 8 sec] 72$^o$C→1 min | 40 | Forward:5'AATGTACCTGTCTGACAGCT 3'<br>Reverse: 5'GTTGGAATACCCTGCCGTGT 3' |

**Table S5: List of Transcription Factors of *GRB2* from MATCH output**

| Factors Name | Position (Strand) | Matrix match score | Sequence (+ strand) |
|---|---|---|---|
| NKx2-5 | 79 (+) | 1.000 | tcAAGTG |
| | 2884 (-) | 1.000 | CACTTga |
| | 7553 (-) | 1.000 | CACTTga |
| | 8983 (+) | 1.000 | tcAAGTG |
| v-Maf | 166 (-) | 0.914 | ctttacttcgTCAGCaagt |
| ER | 478 (-) | 0.964 | tttGGTCAaagttgccttt |
| | 5702 (+) | 0.973 | ggaggttgcagTGACCtga |
| HNF-3beta | 520 (+) | 0.965 | cgcatTATTTtcttt |
| Oct-1 | 843 (-) | 0.917 | attaTTTTCatttca |
| | 883 (+) | 0.939 | tggaatGCAAAtttt |
| | 5448 (-) | 0.988 | ttaaTTTGCatttct |
| | 7142 (+) | 0.955 | gaccatGCAAAttca |
| FOXJ2 | 1608 (+) | 0.948 | gtaACAATattttc |
| HNF-1 | 1786 (+) | 0.852 | gGATAAttagttagctc |
| | 4892 (+) | 0.855 | tGTCAAtagattaccaa |
| Myogenin/NF-1 | 2881 (+) | 0.798 | gatcacttgaggTTGGAgaccagcctggc |
| | 4154 (-) | 0.792 | gaatggcttgaaTCCAGgagacagaggtt |
| | 8072 (+) | 0.781 | cgcctgtgggtgTTGGTagggagctgctc |
| Pax-4 | 3042 (-) | 0.838 | aaggagaatcgCTTGAacgca |
| | 3221 (+) | 0.870 | tgaagTCAGGagtttgagacc |
| | 4027 (+) | 0.876 | cgaggTCAGGagttcaagacc |
| | 4769 (-) | 0.837 | caggagaatcgCTTGAaccca |
| | 8723 (-) | 0.893 | ggctggagacgCTTCActaca |
| CHOP-C/EBPalpha | 3370 (-) | 0.962 | ggagaTTGCAgtg |
| FOXD3 | 3948 (-) | 0.957 | aaataAATAAtt |
| | 8320 (-) | 0.951 | gaataAACATtg |
| COMP1 | 4261 (+) | 0.840 | agaaaaGATTGtcaaggccaggtg |
| | 7734 (-) | 0.814 | actctagcctgggCAATCttagag |
| | 7992 (+) | 0.880 | tttgtgCATTGaccacaagattta |
| AP-1 | 4093 (-) | 0.981 | agttAGTCAgt |
| SREBP-1 | 4335 (+) | 0.995 | gatCACGTgag |
| Evi-1 | 5458 (-) | 0.845 | tTTCTTttcatatgt |
| | 6794 (-) | 0.864 | aTTCTTttcctatca |
| CP2 | 5836 (+) | 0.976 | gctctaTCCAG |
| NF-Y | 7799 (-) | 0.981 | atgATTGGttt |
| Retroviral Poly A | 8663 (-) | 0.973 | cagaacggggctTTATTt |

**Table S6: List of Transcription Factors of *NOX4* from MATCH output**

| Factors Name | Position (Strand) | Matrix match score | Sequence (+ strand) |
|---|---|---|---|
| Oct-1 | 108 (-) | 0.892 | ggaaTTAGCatttgc |
| | 2613 (+) | 0.918 | ggaaatGCTAAtaat |
| | 2648 (+) | 0.923 | atagatGAAAAtta |
| | 3131 (+) | 0.891 | cctgatGCAAAgtgt |
| | 5080 (+) | 0.915 | acttatGAAAAttaa |
| | 6811 (+) | 0.978 | cagaaTTTCAtatc |
| NKx2-5 | 183 (-) | 1.000 | CACTTga |
| | 318 (-) | 1.000 | CACTTga |
| | 3727 (-) | 1.000 | CACTTga |
| | 5776 (+) | 1.000 | tcAAGTG |
| | 8743 (+) | 1.000 | tcAAGTG |
| Pax-4 | 187 (+) | 0.874 | tgaggTCAGGagttgaagacc |
| | 4255 (+) | 0.873 | tgaggTCAGGagttcgagatc |
| | 5783 (-) | 0.901 | ggtccctgaccCCTGAccccc |
| Hand1/E47 | 2420 (-) | 0.970 | aattCCAGAttcagat |
| v-Myb | 3610 (-) | 0.971 | aaCCGTTatc |
| | 6061 (+) | 0.961 | agcAACGGaa |
| Evi-1 | 4148 (-) | 0.874 | tATCTTtttttttct |
| | 8232 (+) | 0.824 | caataaaaaATGATa |
| | 9429 (+) | 0.836 | aaataaaagAGGATa |
| CHOP-C/EBPalpha | 4427 (-) | 0.973 | ggccaTTGCActc |
| HNF-3beta | 4485 (-) | 0.975 | taaatAAATAataac |
| FOXD3 | 4486 (-) | 0.955 | aaataAATAAta |
| YY1 | 5103 (-) | 0.993 | aggagccaagATGGCcgaat |
| Freac-4 | 5552 (+) | 0.983 | cttaggTAAACaaagc |
| Pax-6 | 6354 (-) | 0.884 | atgaaatgaagCGAGAaggga |
| HLF | 7344 (+) | 0.938 | ATTACataat |
| CDP CR3+HD | 7360 (-) | 0.994 | gggATCAAtt |
| COMP1 | 8171 (+) | 0.819 | aaaattGATAGaccactagcaaga |
| | 9620 (-) | 0.843 | cgcatcgccaagtCAATCctaagc |
| CDP CR1 | 8545 (-) | 0.926 | caaTCAATa |
| | 8920 (+) | 0.925 | tATTGAtggg |

**Table S7: Statistical Data for all experiments**

| Figures | Experiments | | Statistical analysis |
|---|---|---|---|
| Figure 1 | **(A)** WB | NOX4 | NOX4 overexpressed by 1.86 fold in AD Brains compared to GAPDH, (n=2), (*p=0.023<0.05) |
| | **(B)** qRT-PCR | NOX4 | NOX4 transcript levels upregulated by 5.06 fold compared to GADH in APP/PS1 mouse brain tissue, (n=2), (*p=0.039<0.05) |
| | **(C)** WB | **(a, b)** Grb2 | Grb2 overexpressed by 5.27 fold in T2D liver compared to GAPDH, (n=2), (*p=0.017<0.05) |
| | | **(a, c)** NOX4 | NOX4 overexpressed by 2.4 fold in T2D liver compared to GAPDH, (n=2), (*p=0.032<0.05) |
| | **(D)** qRT-PCR | Grb2 | Grb2 transcript levels upregulated by 3.01 fold compared to GAPDH in T2D mouse model, (n=3), (*p=0.032<0.05) |
| | **(E)** qRT-PCR | NOX4 | NOX4 transcript levels upregulated by 1.45 fold compared to GAPDH in T2D mouse model, (n=3), (*p=0.032<0.05) |
| | **(F)** WB | Grb2 | **(a) and (ii)** Grb2 overexpressed by 1.22 fold compared to GAPDH in ALK and RYK double knock down condition in SHSY5Y cells, (n=3), (*p=0.027<0.05) |
| | | | **(d) and (iii)** Grb2 overexpressed by 2.34 fold compared to GAPDH in ALK and RYK double knock down condition in HepG2 cells, (n=3), (*p=0.032<0.05) |
| | | NOX4 | **(b) and (iv)** NOX4 overexpressed by 1.35 fold compared to GAPDH in ALK and RYK double knock down condition in SHSY5Y cells, (n=3), (*p=0.039<0.05) |
| | | | **(e) and (v)** Grb2 overexpressed by 2.62 fold compared to GAPDH in ALK and RYK double knock down condition in HepG2 cells, (n=3), (*p=0.022<0.05) |
| | **(G)** qRT-PCR | Grb2 | Grb2 transcript levels upregulated by 4.08 fold compared to GAPDH in ALK and RYK double knock down condition in SHSY5Y cells, (n=3), (*p=0.033<0.05) |
| | | | Grb2 transcript levels upregulated by 5.9 fold compared to GAPDH in ALK and RYK double knock down condition in HepG2 cells, (n=3), (*p=0.009<0.05 |
| | (H) qRT-PCR | NOX4 | NOX4 transcript levels upregulated by 3.02 fold compared to GAPDH in ALK and RYK double knock down condition in SHSY5Y cells, (n=3), (*p=0.023<0.05) |
| | | | NOX4 transcript levels upregulated by 4.13 fold compared to GAPDH in ALK and RYK double knock down condition in HepG2 cells, (n=3), (*p=0.037<0.05) |
| Figure 2 | **(A) (i)** qRT-PCR | NKx2-5 | Nkx2-5 transcript levels downregulated by 3.6 fold compared to GAPDH in AD cell model, (n=3), (*; p=0.0024<0.05) |
| | | PAX4 | PAX4 transcript levels upregulated by 4.19 fold compared to GAPDH in AD cell model, (n=3), |

| | | | |
|---|---|---|---|
| | | | (*;p=0.045<0.05) |
| | | FOXD3 | FOXD3 transcript levels upregulated by 2.89 fold compared to GAPDH in AD cell model, (n=3), (*;p=0.022<0.05) |
| | **(A) (ii)** qRT-PCR | NKx2-5 | Nkx2-5 transcript levels downregulated by 2.59 fold compared to GAPDH in T2D cell model, (n=3), (*;p=0.044<0.05) |
| | | PAX4 | PAX4 transcript levels upregulated by 4.88 fold compared to GAPDH in T2D cell model, (n=3), (*;p=0.024<0.05) |
| | | FOXD3 | FOXD3 transcript levels upregulated by 4.39 fold compared to GAPDH in T2D cell model, (n=3), (*;p=0.046<0.05) |
| | **(B)** WB | PAX4 | **(i) and (ii)** PAX4 overexpressed by 1.8 fold compared to GAPDH in AD patients' whole brain samples, (n=3), (*;p=0.014<0.05) |
| | | | **(i) and (iii)** PAX4 overexpressed by 2 fold compared to GAPDH in T2D patients' whole liver samples, (n=3), (*;p=0.014<0.05) |
| | **(C)** qRT-PCR | **(i)** Grb2 | Grb2 showed no significant alteration (p=0.24>0.05; n=3) in PAX4 knockdown situation in SHSY5Y cells. |
| | | | Grb2 downregulated by 1.52 fold in PAX4 knockdown situation in HepG2 cells, (n=3), (*; p=0.013<0.05) |
| | | **(ii)** NOX4 | NOX4 upregulated by 2.83 fold in PAX4 knockdown situation in SHSY5Y cells, (n=4); (*; p=0.0025<0.05) |
| | | | NOX4 showed no significant alteration (p=0.24>0.05; n=5) in PAX4 knockdown situation in HepG2 cells. |
| | | **(iii)** PAX4 | PAX4 downregulated by 2.99 fold in PAX4 knockdown situation in SHSY5Y cells, (n=3), (*; p=0.012<0.05) |
| | | | PAX4 downregulated by 1.58 fold in PAX4 knockdown situation in HepG2 cells, (n=3), (*; p=0.027<0.05) |
| **Figure 3** | **(A) (i)** qRT-PCR | NKx2-5 | Nkx2-5 transcript levels upregulated by 5.75 fold compared to control siRNAs in ALK and RYK double knock down condition in SHSY5Y cells, (n=4), (*;p=0.016<0.05) |
| | | PAX4 | PAX4 transcript levels upregulated by 5.84 fold compared to control siRNAs in ALK and RYK double knock down condition in SHSY5Y cells, (n=3), (**; p=0.00015<0.001) |

| | | | |
|---|---|---|---|
| | | FOXD3 | FOXD3 transcript levels upregulated by 3.13 fold compared to control siRNAs in ALK and RYK double knock down condition in SHSY5Y cells, (n=4),<br>(*; p=0.045<0.05) |
| | **(A) (ii)** qRT-PCR | NKx2-5 | Nkx2-5 showed no significant alteration of transcript level (p=0.24>0.05; n=5) in ALK and RYK double knock down condition in HepG2 cells, (n=3), (N.S.; p=0.15>0.05) |
| | | PAX4 | PAX4 transcript levels upregulated by 4.81 fold compared to control siRNAs in ALK and RYK double knock down condition in HepG2 cells, (n=3),<br>(*; p=0.003<0.05) |
| | | FOXD3 | FOXD3 transcript levels upregulated by 2.46 fold compared to control siRNAs in ALK and RYK double knock down condition in HepG2 cells, (n=4),<br>(*; p=0.041<0.05) |
| | **(B)** WB | PAX4 | **(i) and (ii)** PAX4 overexpressed by 1.65 fold compared to control siRNAs in ALK and RYK double knock down condition in SHSY5Y cells, (n=3), (*;p=0.026<0.05) |
| | | | **(i) and (iii)** PAX4 overexpressed by 1.98 fold compared to control siRNAs in ALK and RYK double knock down condition in HepG2 cells, (n=3), (*;p=0.004<0.05) |
| | (C) ChIP qRT-PCR | Grb2 | (i) PAX4 binding significantly increase the Grb2 expression levels by 27.49 fold in AD cell model, (n=3), (*; p=0.005<0.05) |
| | | | (ii) PAX4 binding significantly increase the Grb2 expression levels by 18.33 fold in T2D cell model, (n=3), (*;p=0.019<0.05) |
| | | NOX4 | (i) PAX4 binding showed no significant alteration of NOX4 transcript level in AD cell model. (N.S.; p=0.18>0.05) |
| | | | (ii) PAX4 binding showed no significant alteration of NOX4 transcript level in T2D cell model. (N.S.; p=0.15>0.05) |
| **Figure 4** | (A) qRT-PCR | ARX | ARX transcript level upregulated by 3.35 fold in PAX4 knockdown condition compared to mock siRNAs in SHSY5Y cells. (n=5); (*p=0.016<0.05). |
| | | | ARX transcript level upregulated by 3.13 fold in PAX4 knockdown condition compared to mock siRNAs in HepG2 cells. (n=8); (*p=0.0012<0.05). |
| | (B) qRT-PCR | ARX | ARX transcript level downregulated by 2.6 fold in AD cell model (AICD+Aβ)compared to control condition (GFP+DMSO). (n=3); (*p=0.04<0.05). |
| | | | ARX transcript level downregulated by 3.7 fold in T2D cell model (+PA+Amylin+Insulin) compared to control condition (-PA-Amylin+Insulin). (n=4); (**p=0.0003<0.001). |

| | | | |
|---|---|---|---|
| | | | ARX transcript level downregulated by 3 fold in ALK and RYK double knockdown condition compared to mock siRNAs in SHSY5Y cells. (n=3); *p=0.03<0.05). |
| | | | ARX transcript level downregulated by 3.7 fold in ALK and RYK double knockdown condition compared to mock siRNAs in HepG2 cells. (n=3); *p=0.02<0.05). |
| | (C) WB | β-Catenin | **(i) and (iii)** β-Catenin downregulated by 3.5 fold in AD cell model, (n=3), (**p=0.0006<0.001) |
| | | | **(ii) and (iii)** β-Catenin downregulated by 1.5 fold in T2D cell model, (n=3), (**p=0.0006<0.001) |
| | (D) WB | β-Catenin | **(i) and (iii)** β-Catenin downregulated by 1.15 fold in ALK and RYK double knockdown condition in SHSY5Y cells, (n=3), (*p=0.015<0.05) |
| | | | **(ii) and (iii)** β-Catenin downregulated by 6.5 fold in ALK and RYK double knockdown condition in HepG2 cells, (n=2), (*p=0.013<0.05) |
| **Figure 5** | (A) WB | (a) (i), (b) α-Tubulin | α-Tubulin showed no significant alteration in T2D liver lysate (n=2), (p=0.19>0.05) |
| | | (a) (ii), (b) Vimentin | Vimentin downregulated by 1.83 fold in T2D liver lysate compared to WT (n=2), (*p=0.011<0.05) |
| | | (a) (iii), (b) α-SMA | α-SMA downregulated by 1.68 fold in T2D liver lysate compared to WT (n=2), (*p=0.044<0.05) |
| | | (a) (iv), (b) Stathmin1 | Stathmin1 downregulated by 1.95 fold in T2D liver lysate compared to WT (n=2), (*p=0.024<0.05) |
| | (B) qRT-PCR | α-Tubulin | α-Tubulin downregulated by 5.3 fold in T2D mouse model compared to WT (n=3), (*p=0.0013<0.05) |
| | | Vimentin | Vimentin downregulated by 6.49 fold in T2D mouse model compared to WT (n=3), (*p=0.017<0.05) |
| | | α-SMA | α-SMA downregulated by 2 fold in T2D mouse model compared to WT (n=3), (*p=0.028<0.05) |
| | | Stathmin1 | Stathmin1 downregulated by 2.9 fold in T2D mouse model compared to WT (n=3), (*p=0.009<0.05) |
| | (C) WB | (a) (i), (b) α-Tubulin | α-Tubulin downregulated by 2.7 fold in T2D cell model compared to WT (n=2), (**; p=0.00017<0.001) |
| | | (a) (ii), (b) Vimentin | Vimentin downregulated by 1.98 fold in T2D cell model compared to WT (n=2), (*;p=0.037<0.05) |
| | | (a) (iii), (b) α-SMA | α-SMA downregulated by 1.27 fold in T2D cell model compared to WT (n=4), (*;p=0.004<0.05) |
| | | (a) (iv), (b) Stathmin1 | Stathmin1 downregulated by 1.97 fold in T2D cell model compared to WT (n=4), (***;p<0.0001) |
| | (D) WB | (i), (ii) α-Tubulin | α-Tubulin level downregulated by 2.64 fold in ALK and RYK double knockdown condition compared to mock siRNAs in SHSY5Y cells. (n=3), (*; p=0.0507≈0.05) |
| | | | α-Tubulin level downregulated by 1.42 fold in ALK and RYK double knockdown condition |

| | | | |
|---|---|---|---|
| | | | compared to mock siRNAs in HepG2 cells. (n=3), (*;p=0.019<0.05) |
| | (E) qRT-PCR | α-Tubulin | α-Tubulin transcript level downregulated by 2.3 fold in ALK and RYK double knockdown condition compared to mock siRNAs in SHSY5Y cells. (n=3), (*; p=0.03<0.05) |
| | | | α-Tubulin transcript level downregulated by 1.5 fold in ALK and RYK double knockdown condition compared to mock siRNAs in HepG2 cells. (n=3), (*; p=0.021<0.05) |
| | (F) qRT-PCR | α-Tubulin | α-Tubulin level downregulated by 1.58 fold in the T2D cell model. (n=3), (*; p=0.018<0.05) |
| | | | α-Tubulin level upregulated by 1.24 fold in the T2D cell model along with Grb2 overexpression. (n=3), (*; p=0.018<0.05) |
| | | Vimentin | Vimentin level downregulated by 2.12 fold in the T2D cell model. (n=3), (*;p=0.024<0.05) |
| | | | Vimentin level upregulated by 1.15 fold in the T2D cell model along with Grb2 overexpression. (n=3), (*; p=0.037<0.05) |
| | | α-SMA | α-SMA level downregulated by 2.2 fold in the T2D cell model. (n=3), (*;p=0.027<0.05) |
| | | | α-SMA level upregulated by 2.76 fold in the T2D cell model along with Grb2 overexpression. (n=3), (*; p=0.023<0.05) |
| | | Stathmin1 | Stathmin1 level downregulated by 1.75 fold in the T2D cell model. (n=3), (*;p=0.002<0.05) |
| | | | Stathmin1 level upregulated by 1.07 fold in the T2D cell model along with Grb2 overexpression. (n=3), (*; p=0.021<0.05) |
| **Figure 6** | (A) WB | (i), (b) RhoA | RhoA downregulated by 2.13 fold in the T2D cell model compared to WT (n=3), (*; p=0.012<0.05) |
| | | | RhoA downregulated by 1.22 fold in the T2D cell model along with the overexpressed Grb2 compared to WT (n=3), (*;p=0.025<0.05) |
| | | (ii), (b) Rac1 | Rac1 downregulated by 1.63 fold in the T2D cell model compared to WT (n=3), (*; p=0.0051<0.05) |
| | | | Rac1 downregulated by 1.26 fold in the T2D cell model along with the overexpressed Grb2 compared to WT (n=3) (*; p=0.0065<0.05) |
| | | (iii), (b) Cdc42 | Cdc42 upregulated by 1.36 fold in the T2D cell model compared to WT (n=3), (*; p=0.013<0.05) |
| | | | Cdc42 downregulated by 2.05 fold in the T2D cell model along with the overexpressed Grb2 compared to WT (n=3) (*; p=0.02<0.05) |
| | (B) WB | (a) (i), (b) (i) Cofilin | The Cofilin activation level downregulated by 1.43 fold in the T2D cell model compared WT(n=3) (*; p = 0.011<0.05) |
| | | | The Cofilin activation level showed no significant alteration in the T2D cell model along with overexpressed Grb2 compared to WT (n=3) (N.S; p=0.99) |
| | | (a) (ii), (b) (ii) NOX4 | NOX4 upregulated by 1.31 fold in the T2D cell model compared to WT (n=3) (*; p = 0.042<0.05) |

| | | | |
|---|---|---|---|
| | | | The NOX4 level downregulated by 1.26 fold in the T2D cell model along with overexpressed Grb2 compared to WT (n=3) (*; p=0.01<0.05) |
| | (C) WB | (a) (i), (b) Gsk3β | Gsk3β activity levels upregulated by 1.7 fold in the T2D cell model compared to WT (n=3) (*; p=0.01>0.05) |
| | | (a) (ii), (b) AKT | AKT activity levels downregulated by 1.54 fold in the T2D cell model compared to WT (n=3) (**; p=0.00005<0.001) |
| **Figure 7** | (A) Co-IP | (a) NOX4 | NOX4 and Grb2 interaction upregulated by 1.35 fold in the T2D cell model compared to WT (n=3) (*; p=0.0197<0.05) |
| | | (b) NOX4 | NOX4 and Grb2 interaction upregulated by 2.87 fold in the T2D mouse model compared to WT. |
| | (B), (C) ROS actiivity | CMH2-DCFDA | The CMH2-DCFDA absorbance upregulated by 1.32 fold in T2D cell model compared to WT (n=3) (*; p=0.0346<0.05) |
| | | | The CMH2-DCFDA absorbance upregulated by 3.57 fold in T2D cell model along with overexpressed Grb2 compared to WT (n=3) (*; p=0.005<0.05) |
